# Physiological Noise Correction in Brainstem Imaging: An Empirical Evaluation of fMRI Data-Driven and Peripheral Physiological Recording-Driven Methods

**DOI:** 10.1101/2023.02.22.529506

**Authors:** Martin Krentz, Rayyan Tutunji, Nikos Kogias, Hariharan Murali Mahadevan, Zala C. Reppmann, Florian Krause, Erno J. Hermans

## Abstract

Physiological noise substantially affects fMRI data quality, particularly in areas near fluid-filled cavities and arteries such as the brainstem. Peripheral physiological signals can be recorded alongside fMRI and incorporated into general linear models to remove variance associated with cardiac and respiratory cycles using methods such as Retrospective Image Correction (RETROICOR). In contrast, data-driven noise reduction methods such as anatomical component correction (aCompCor) and independent component analysis based automatic removal of motion artifacts (ICA-AROMA) do not rely on additional peripheral physiological recordings. These methods have shown efficacy in correcting for motion, scanner as well as physiological artifacts. This raises the question of whether logistically demanding peripheral recordings offer added value. We therefore used nuisance regression methods based on peripherally recorded physiology (RETROICOR, heart rate, respiratory volume) as well as fMRI data itself (ICA-AROMA, aCompCor) to account for noise in a resting-state data set. Subsequently, we compared variance explained by the respective methods and improvements in temporal signal-to-noise ratio throughout different regions of interest in the brain. Consistent with prior research, RETROICOR, heart rate and respiratory volume explain significant variance throughout the brain with peaks around areas of strong cardiac pulsations. ICA-AROMA and aCompCor account for a substantial proportion of the variance explained by the methods using peripheral physiology. Nonetheless, the methods based on peripheral physiological recordings retain unique explanatory power. Analysis revealed a pattern of unreliability of ICA-AROMA to consistently identify and remove physiological noise across recordings in each participant, which is compensated by RETROICOR, heart rate and respiratory volume. This unreliability partially results from misclassifications in the noise selection models of ICA-AROMA. Our results suggest that it is advisable to additionally apply cleaning based on peripheral physiological recordings, especially when assessing inter-individual differences and areas with regionally high levels of physiological noise.

## 1. Introduction

### 1.1. Rising interest in brainstem imaging

Functional imaging of brainstem regions has become increasingly important for research in a variety of neuroscience sub-fields, ranging from investigations of task-related activity in brainstem nuclei to functional connectivity with higher-order brain networks (Chu et al., 2021; Colizoli et al., 2022; Henderson & Macefield, 2013). For instance, the locus coeruleus (LC) has seen a surge of interest, because of the crucial role it plays in a range of processes as the primary source of cortical norepinephrine (Elman et al., 2017; Schwarz & Luo, 2015). Despite the interest, it is largely unexplored regarding its functional role in neuroimaging studies due to its high susceptibility to noise interference, primarily physiological noise. While availability of higher-field strength MRI is growing and allowing for higher resolution imaging and progress in usage of novel sequences to structurally identify different brainstem nuclei (Singh et al., 2022), the functional acquisition of their respective activity remains notoriously difficult. Over the past two decades a variety of tools have been proposed to reduce the impact of physiological noise on fMRI data quality (Brooks et al., 2013; Caballero-Gaudes & Reynolds, 2017; Murphy et al., 2013). However, as the available solutions differ in accessibility, effort and ease of use, questions about their relative theoretical benefits have to be weighed against practical considerations. With the increasing emphasis on reproducibility and ease of access, it is important to assess whether standard processing pipelines such as fMRIPrep (Esteban, Markiewicz, et al., 2018), which streamline pre-processing steps and incorporate noise correction methods like aCompCor (anatomical Component Based Noise Correction; Behzadi et al., 2007) and allow accessible subsequent computation of methods such as ICA-AROMA denoising (Automatic Removal of Motion Artifacts; Pruim et al., 2015), can serve as effective substitutes for more labor- and expertise-intensive approaches that rely on peripheral physiological recordings (Birn et al., 2006; Glover et al., 2000), even for imaging of brainstem areas.

### 1.2. Sources of physiological noise

Classically, physiological noise describes fMRI signal variation attributed to physiological processes, such as cardiac and respiratory activity, which are unrelated to neural signal of interest (i.e. task or rest BOLD effects). These undesired signal variations are diverse in their temporal and spatial characteristics, as well as their discernability and acting mechanisms.

#### 1.2.1. Non-BOLD effects of physiological noise

One of the most prominent components, often associated with cyclic cardiac artifacts, are cardiac pulsations, leading to voxel displacement and partial-volume effects prominently in areas proximal to large arteries or CSF areas such as 3^rd^ or 4^th^ ventricle (Dagli et al., 1999; Glover et al., 2000; T. T. Liu, 2016). Similarly, cyclic respiratory activity shows indirect BOLD effects through breathing induced organ movement and susceptibility changes, resulting in magnetic field change (B0) as well as modulation of intravoxel dephasing (Raj et al., 2001) and venous pressure (Windischberger et al., 2002). In contrast to cardiac cyclic artifacts, the localization of respiratory effects appears stronger along the edges of the brain and ventricles (Van de Moortele et al., 2002; Windischberger et al., 2002).

#### 1.2.2. BOLD effects of physiological noise

Beyond effects on the quality of data acquisition itself and limitations in accuracy of the BOLD assessment, low frequency physiological processes such as heart rate frequency changes (Shmueli et al., 2007) or changes in CO2 as a potent vasodilator (Birn et al., 2006, 2008; Wise et al., 2004) can directly affect the BOLD signal. Furthermore, an interdependency between cardiac and respiratory activity exists, reflecting itself in respiratory cycle dependent heart rate changes (sinus arrhythmia, Berntson et al., 1993) as well as CSF flow, hypothesized to be largely driven by changes in negative intrathoracic venous pressure as a consequence of inhalation (Dreha-Kulaczewski et al., 2015; Kollmeier et al., 2022).

Since measured BOLD is dependent on cerebral blood flow (CBF) as well as cerebral blood volume (CBV), heart rate or respiration induced blood pressure and CO2 changes directly affect the derived signal (Das et al., 2021; T. T. Liu, 2016). These changes in difference to artifacts created by the cyclic higher frequency cardiac (∼ 1.3Hz) and respiratory (∼ 0.5Hz) dynamics, act in low frequency bands (∼ 0.1Hz) and thus are especially detrimental for methodology such as resting-state functional connectivity which relies on synchronized low-frequency changes in activity (Kelly et al., 2012; Zuo et al., 2019).

While the described physiological noise has a varying impact on all BOLD-based brain functional assessments, brainstem regions are specifically affected due to their aforementioned high susceptibility to arterial and CSF pulsatility and respiratory magnetization effects (Brooks et al., 2013; Krüger & Glover, 2001; T. T. Liu, 2016). This difficulty is amplified by the general nature of assessed brainstem nuclei due to their small size compared to cortical gray matter regions, proximity to other brainstem regions with distinct functionality and direct edge proximity to areas of non-interest such as the 4^th^ ventricle. Some of these downsides could be compensated with high-resolution fMRI assessments using high field-strength (i.e. 7T) and improved receiver coil positioning and sensitivity. However, while high-field fMRI delivers on higher resolution and spatial discernability, it is disproportionally affected by physiological noise and thus makes further compensation of physiological noise is paramount (Hutton et al., 2011; Krüger & Glover, 2001; Triantafyllou et al., 2005).

### 1.3. Correcting for physiological noise

Over the last decade a wide range of improvements has been made in regards to tackling noise in resting-state fMRI analysis. Not only with respect to the quality of noise correction, but also its ease in applicability. With a push for reproducible science and lower barrier of access, software packages such as ‘fMRIPrep’ (Esteban, Markiewicz, et al., 2018) have been growing in popularity; making access to certain noise correction pipelines less dependent on high levels of localized expertise. Due to the complexity of physiological noise dynamics, a wide variety of methods have been proposed to tackle different aspects of the problem at different stages of acquisition and analysis (Caballero-Gaudes & Reynolds, 2017; Murphy et al., 2013).

#### 1.3.1. Using peripheral recordings for physiology clean-up

Most prominent approaches of physiological noise correction focus on the post-hoc removal of physiologically induced artifacts. One possibility is to account for physiological noise by acquiring cardiac and respiratory measures during fMRI assessments. The most common approach – and often treated as the gold standard in physiological noise correction – is RETROICOR (Glover et al., 2000), a Fourier series of cardiac and respiratory signal to model the phase of the respective cycles. This phase information is typically used for nuisance regression by being included into planned general linear models. Inclusion of phase information specifically aims to reduce effects of cardiac pulsatility and motion-like respiratory effects onto the main magnetic field. In line with the higher susceptibility to physiological noise, areas in, or proximal to, the brainstem – especially areas surrounding main cerebral blood flow i.e. Circle of Willis – have shown the biggest improvement, despite overall efficacy in reducing physiological noise across the cortex (Glover et al., 2000; Van Buuren et al., 2009). In addition to RETROICORs correction of respiratory and cardiac phase-related signal changes, other effective measures derived from physiological recordings aim at accounting for the variation of respiration and cardiac activity over time. Respiratory volume per unit time (RVT), which encompasses both respiratory rate and depth changes time locked to a volume, allows to remove physiological noise associated to changes in breathing patterns across the brain (Birn et al., 2006). While inclusion of time lagged cardiac rate as a means to capture delayed cardiac frequency-related signal change further removes mostly spatially unspecific physiological noise in both grey and white matter in resting state fMRI (Chang et al., 2009; Shmueli et al., 2007). While only suffering from the minimal downside of loss of temporal degrees of freedom (tDOF), this combination of approaches puts extra strain onto the logistics of an experimental set-up. It necessitates the application of extra equipment leading to increased preparation time and sources of potential malfunctions or experimental inconsistencies.

#### 1.3.2. Data-driven approaches of physiological clean-up

An alternative approach for noise removal is to employ temporo-spatial decomposition of the data using independent or principal component analysis. This approach has been used to great success in identifying dynamics of functional-brain networks in resting-state fMRI (Beckmann et al., 2005; Beckmann & Smith, 2004). Since it can be used for identification of systematic temporo-spatial signal patterns, it similarly can be used to identify components of non-interest, such as head motion- and scanner artifacts, as well as some aspects of physiological noise (Pruim, Mennes, van Rooij, et al., 2015). While powerful, the main difficulty of using an ICA-approach is the necessary prior knowledge for correct classification of independent components as noise. In an effort to create an automated component selection, independent of rater expertise, Pruim and colleagues (2015) suggest an automatic tool for removal of noise artifacts named ICA-AROMA. Different to other solutions, such as ICA-FIX (Salimi-Khorshidi et al., 2014) which needs a training data-set to run a classifier and subsequent component selection, ICA-AROMA uses a pre-trained classifier based on three decision criteria for each component: (1) whether it contains high-frequency content unlikely to be brain signal, (2) whether its spatial layout shows high overlap with brain edges and (3) whether its spatial layout strongly overlaps with CSF regions. While this approach was originally designed to account for head motion and scanner artifacts, ICA-AROMA assesses features that are equally crucial in detecting physiological noise. The approach of using shrunk brain masks to calculate edge fraction allows for possible identification of respiratory bulk head motion or effects visible in venous drainage in the sagittal sinus. More importantly, however, the CSF fraction approach classifies based on IC component spatial distribution along edges of CSF within the brain – typically regions most affected by cardiac pulsatility. Thus, while ICA-AROMA does not aim for physiological noise correction, its mechanisms make detection likely. ICA-AROMA can be furthermore combined with aCompCor, a method utilizing principal components of white matter- and CSF masks as nuisance variables (Behzadi et al., 2007), which is commonly used for physiological noise correction. Similar to ICA-AROMA, it can be derived post-hoc from pre-processed data and has shown promise in approaching the efficacy of methods based on peripheral physiology such as RETROICOR (Behzadi et al., 2007; Muschelli et al., 2014). The joint use of these pipelines makes it a ready-made approach of correcting for different types of noise sources, including physiological noise.

### 1.4. Commonalities and differences between noise correction methods

In this study we assess whether more logistically costly methods derived directly from peripheral physiological recordings, such as RETROICOR, heart rate (HR) and respiratory volume per unit time (RVT), have added benefit on top of fMRI data-driven methods such as ICA-AROMA and aCompCor. Particularly, we investigate if this is the case in regions highly affected by physiological noise such as the brainstem. To test group-level differences we assess the shared and unique contributions to improvement of temporal signal to noise ratio (tSNR) as a consequence of the different correction methods for separate regions of interest. The regions are chosen for different levels of specificity (whole-brain, gray matter, brainstem and locus coeruleus) in regard to proximity and likelihood of high physiological noise. Furthermore, by testing the contribution of the respective approaches to image variance, we hope to identify common components of noise signal and their consistency across a variety of participants. Using the information about spatial patterns gained from cardiac and respiratory regressors, we investigate the efficacy of ICA-AROMA in regard to its inter-individual consistency and stability.

## 2. Methods

### 2.1. Participants

Thirty healthy volunteers (15 females, 15 males) between 18 and 55 years were recruited for this experiment via the ‘Radboud Research Participation System’ and compensated 60€ for their participation. Participants were right-handed, healthy, and without a history of neurological or psychiatric illness. Additionally, MRI contraindications, such as ferromagnetic implants or a history of claustrophobia, were an exclusion criterion. Before completion of the experimental sessions, participants received instructions and information about the experimental procedures. The data used in this article is part of a larger study involving multiple sessions and manipulations of no relevance for the here described approach. Two participants did not complete all sessions and scans and were thus excluded from further analysis. Due to a technical problem with the heart-rate acquisition, one participant had to be excluded from the current analysis. Thus, a total of 27 subjects were used for reported analyses (12 females, 15 males; mean age = 21.45, SD = 3.6). An error in locus coeruleus imaging led to a missing mask for one subject, reducing the number of subjects included for the LC ROI analysis of LC to 26. The study was approved by the local ethics committee, and participants gave their written informed consent before the procedure.

### 2.2. Design

The study consisted of three separate sessions including behavioral measures not relevant to this study, as well as structural and functional MRI acquisitions. In an initial session, structural images were acquired including a T1-weighted image as well as a combination of turbo spin-echo (TSE) sequences for identification of the Locus Coeruleus (Sasaki et al., 2006). Sessions two and three containing functional resting state runs were identical, except for a counter-balanced stress-induction procedure administered before the start of scanning without further relevance for the current analysis.

#### 2.2.1. Procedure

Before entering the MR scanner, participants were attached to all physiological recording devices while seated on the scanner bed. Subsequently, participants were placed into the head coil with firm foam cushioning around their head to reduce head movement. Additionally, a piece of tape was placed over their forehead to further reduce motion (Krause et al., 2019).

#### 2.2.2. MRI data acquisition

All MR images were recorded on a Siemens Prisma-Fit 3T MR System (Siemens, Erlangen, Germany) using a 32-channel receiver head coil. Structural images were acquired using a T1-weighted sequence with using a 3D resolution of 0.8x0.8x0.8 mm (repetition time = 2.2s, echo time = 26.4ms, flip angle = 11, field of view 256x256mm, slices = 320. Functional images were acquired with a BOLD contrast sensitive EPI sequence using a multiband acceleration factor of 3. For higher sensitivity in brainstem resolution, we chose a slice thickness of 1.5 mm (no gap) with an in-plane resolution of 1.5x1.5 mm (repetition time = 2020ms, echo time = 28.4ms, flip angle = 80, field of view = 210x210 mm, slices = 87) and a generalized auto calibrating partial parallel acquisition (GRAPPA) factor of 2 for a total number of 240 volumes. For structural assessment of the locus coeruleus (LC) and delineation as a functional mask, we used a T1-weighted turbo spin-echo sequence with slice thickness of 3mm and an in-plane resolution of 0.24x0.24 mm aligned perpendicular to the brainstem (repetition time = 1870ms, echo time = 12ms, flip angle = 120, slices = 16, field of view = 220x200mm) (Sasaki et al., 2006). Detailed processing mechanisms for mask creation are presented in Appendix A.

### 2.3. MRI data preprocessing

#### 2.3.1. Anatomical data preprocessing

Results included in this manuscript come from preprocessing performed using fMRIPrep 23.0.2 (Esteban et al., 2019; Esteban, Blair, et al., 2018; RRID:SCR_016216), which is based on Nipype 1.8.6 (Gorgolewski et al., 2011, 2018; RRID:SCR_002502). Further processing was done using custom python scripts. A B0 nonuniformity map (or fieldmap) was estimated from the phase-drift map(s) measure with two consecutive GRE (gradient-recalled echo) acquisitions. The corresponding phase-map(s) were phase-unwrapped with prelude (FSL 6.0.5.1:57b01774). The T1-weighted (T1w) image was corrected for intensity non-uniformity (INU) with N4BiasFieldCorrection (Tustison et al., 2010), distributed with ANTs 2.2.0 (Avants et al., 2008; RRID:SCR_004757), and used as T1w-reference throughout the workflow. The T1w-reference was then skull-stripped with a Nipype implementation of the antsBrainExtraction.sh workflow (from ANTs), using OASIS30ANTs as target template. Brain tissue segmentation of cerebrospinal fluid (CSF), white-matter (WM) and gray-matter (GM) was performed on the brain-extracted T1w using fast (FSL 6.0.5.1:57b01774, RRID:SCR_002823, Zhang et al., 2001). Brain surfaces were reconstructed using recon-all (FreeSurfer 7.3.2, RRID:SCR_001847, Dale et al., 1999), and the brain mask estimated previously was refined with a custom variation of the method to reconcile ANTs-derived and FreeSurfer-derived segmentations of the cortical gray-matter of Mindboggle (RRID:SCR_002438, Klein et al., 2017). Volume-based spatial normalization to two standard spaces (MNI152NLin6Asym, MNI152NLin2009cAsym) was performed through nonlinear registration with antsRegistration (ANTs 2.3.3), using brain-extracted versions of both T1w reference and the T1w template. The following templates were selected for spatial normalization and accessed with TemplateFlow (23.0.0, Ciric et al., 2022): FSL’s MNI ICBM 152 non-linear 6th Generation Asymmetric Average Brain Stereotaxic Registration Model [Evans et al., 2012; RRID:SCR_002823; TemplateFlow ID: MNI152NLin6Asym], ICBM 152 Nonlinear Asymmetrical template version 2009c [Fonov et al., 2009, RRID:SCR_008796; TemplateFlow ID: MNI152NLin2009cAsym]. For LC imaging adjustments were made to the MR sequence parameters used by Sasaki et al. (2006) based on in-house testing results showing a maximization of contrast. An in-house pipeline was used (Supplement A) to delineate a usable LC mask for functional signal extraction on an individual level.

#### 2.3.2. Functional image processing

For each individual resting state run the following preprocessing was performed. First, a reference volume and its skull-stripped version were generated using a custom methodology of fMRIPrep. Head-motion parameters with respect to the BOLD reference (transformation matrices, and six corresponding rotation and translation parameters) are estimated before any spatiotemporal filtering using mcflirt (FSL 6.0.5.1:57b01774, Jenkinson et al., 2002). The estimated fieldmap was then aligned with rigid-registration to the target EPI (echo-planar imaging) reference run. The field coefficients were mapped on to the reference EPI using the transform. The BOLD reference was then co-registered to the T1w reference using bbregister (FreeSurfer) which implements boundary-based registration (Greve & Fischl, 2009). Co-registration was configured with six degrees of freedom. Several confounding time-series were calculated based on the preprocessed BOLD: framewise displacement (FD), DVARS and three region-wise global signals. FD was computed using two formulations following Power (absolute sum of relative motions, Power et al., 2014) and Jenkinson (relative root mean square displacement between affines, Jenkinson et al., 2002). FD and DVARS are calculated for each functional run, both using their implementations in Nipype (following the definitions by Power et al. 2014). The three global signals are extracted within the CSF, the WM, and the whole-brain masks. Additionally, a set of physiological regressors were extracted to allow for component-based noise correction (CompCor, Behzadi et al. 2007). Principal components are estimated after high-pass filtering the preprocessed BOLD time-series (using a discrete cosine filter with 128s cut-off) for the two CompCor variants: temporal (tCompCor) and anatomical (aCompCor). tCompCor components are then calculated from the top 2% variable voxels within the brain mask. For aCompCor, three probabilistic masks (CSF, WM and combined CSF+WM) are generated in anatomical space. The implementation differs from that of Behzadi et al. in that instead of eroding the masks by 2 pixels on BOLD space, a mask of pixels that likely contain a volume fraction of GM is subtracted from the aCompCor masks. This mask is obtained by dilating a GM mask extracted from the FreeSurfer’s aseg segmentation, and it ensures components are not extracted from voxels containing a minimal fraction of GM. Finally, these masks are resampled into BOLD space and binarized by thresholding at 0.99 (as in the original implementation). Components are also calculated separately within the WM and CSF masks. For each CompCor decomposition, the k components with the largest singular values are retained, such that the retained components’ time series are sufficient to explain 50 percent of variance across the nuisance mask (CSF, WM, combined, or temporal). The remaining components are dropped from consideration. The head-motion estimates calculated in the correction step were also placed within the corresponding confounds file. The confound time series derived from head motion estimates and global signals were expanded with the inclusion of temporal derivatives and quadratic terms for each (Satterthwaite et al., 2013). Frames that exceeded a threshold of 0.5 mm FD or 1.5 standardized DVARS were annotated as motion outliers. Additional nuisance timeseries are calculated by means of principal components analysis of the signal found within a thin band (crown) of voxels around the edge of the brain, as proposed by Patriat et al. (2017). The BOLD time-series were resampled into standard space, generating a preprocessed BOLD run in MNI152NLin6Asym space. Automatic removal of motion artifacts using independent component analysis (ICA-AROMA, Pruim, Mennes, van Rooij, et al., 2015) was performed on the preprocessed BOLD on MNI space time-series after removal of non-steady state volumes and spatial smoothing with an isotropic, Gaussian kernel of 6mm FWHM (full-width half-maximum). The “aggressive” noise-regressors were collected and placed in the corresponding confounds file. All resamplings can be performed with a single interpolation step by composing all the pertinent transformations (i.e. head-motion transform matrices, susceptibility distortion correction when available, and co-registrations to anatomical and output spaces). Gridded (volumetric) resamplings were performed using antsApplyTransforms (ANTs), configured with Lanczos interpolation to minimize the smoothing effects of other kernels (Lanczos, 1964). Non-gridded (surface) resamplings were performed using mri_vol2surf (FreeSurfer). Components classified as noise after running MELODIC-ICA (FSL), were concatenated per subject to create a regressors matrix of AROMA noise regressors. The AROMA noise regressors are of variable length, as both the number of ICA components as well as classifications as noise varies across participants as a function of motion and other noise sources. These concatenated regressors will further be called ‘AROMA regressors’. Furthermore, aCompCor components were extracted from the nuisance regressor file created by fmriprep. For the analysis the first 6 PCA components derived from white matter and CSF masks as implemented in aCompCor were used. These 6 components were concatenated and are further called ‘aCompCor regressors’.

#### 2.3.3. Physiological data acquisition

Physiological data were recorded using a finger pulse sensor placed on the index finger (Brain Products, Gilching, Germany) connected to a BrainAmp ExG MR (Brain Products, Gilching, Germany). An abdominal respiration belt (attached to a pneumatic sensor, Brain Products; Gilching, Germany) was used to assess respiratory activity. Acquired physiological data were subsequently processed using an in-house tool for manual noise rejection, peak detection, and quality control (https://github.com/can-lab/hera). The outputs were a corrected time series for the cardiac acquisition. Respiratory data was transformed using another in house tool and concurrently cleaned for possible artifacts (https://github.com/can-lab/brainampconverter).

Subsequently, the processed physiological data was input into an inhouse MATLAB-based implementation of RETROICOR (v2.1; https://github.com/can-lab/RETROICORplus) combining trigger information and physiology to create multiple regressors modeling the phase of both cardiac and respiratory rate (Glover et al., 2000). The final model was reduced to include third order cardiac phase regressors and fourth order respiratory phase regressors, as well as a multiplication term of the first cardiac and respiratory phase regressors, which was calculated to account for possible interaction effects (Harvey et al., 2008). This resulted in a total of 18 regressors further called ‘RETROICOR regressors’. For a detailed view of recording data quality see Appendix B. Additionally, two additional types of regressors were included: (1) A collection of three regressors modeling running heart rate frequency, calculated per volume and time delayed by one and two volumes respectively, further called ‘HR regressors’. (2) A set of two regressors with one containing the product of respiratory frequency and amplitude, averaged per volume within a 9 second window and a second regressor using a lag of one volume to model time shifted dynamics, further called ‘RVT regressors’.

### 2.4. Pipeline for correctional method comparison

#### 2.4.1. General linear models integrating physiological noise correction

Regressors resulting from all described correction methods, are typically used as nuisance variables in general linear models (GLMs) to remove variance attributed to various sources of noise from signal of interest. To directly assess the impact of the different noise correction procedures on an individual level we chose to compute GLMs treating these commonly used nuisance variables as variables of interest to directly assess the variance explained by the respective procedures. Furthermore, to assess how the nuisance regressors based on recorded physiological data compare to the fMRI data-derived regressors of AROMA and aCompCor, combinatory GLMs were created. In total, 9 GLMs were computed using Nilearn (https://github.com/nilearn; Abraham et al., 2014). The contrasts computed for all GLMs were z-transformed and FDR-thresholded (p < .05) to ensure the detection of spatial patterns. A GLM was computed for each set of nuisance regressors respectively, without further addition of motion parameters.

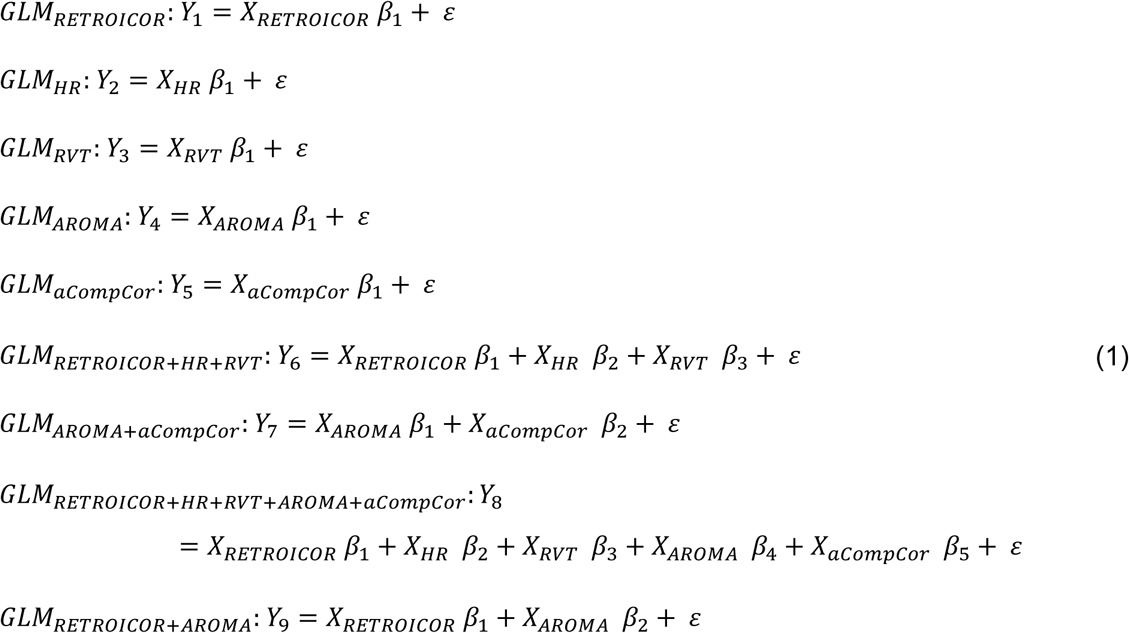

Consequently, an F-contrast was computed across all regressors for each method respectively. Furthermore, the combined methods based on recorded physiological data as well as fMRI data-derived methods were tested together to be used for later TSNR comparison (GLM_RETROICOR+HR+RVT_/GLM_AROMA+aCompCor_). A combinatory GLM was run including RETROICOR, HR, RVT, aCompCor, and AROMA regressors to assess unique contribution of methods based on physiological recordings beyond a combination of both data-driven methods. An F-contrast was computed across all RETROICOR, HR and RVT regressors. To assess the unique contribution of RETROICOR to ICA-AROMA we computed a GLM using RETROICOR as well as AROMA regressors (GLM_RETROICOR+AROMA_). While not used for the main analysis, this GLM is used to identify ICA-AROMA misclassification as described in section 2.5. All models were run without further nuisance regression. No second-level statistics were calculated based on these models, as (1) the main focus was an individual assessment of stability and individual impact of the chosen respective noise correction methods (2) limitations of using F-contrasts for second-level statistics in this context due to the lack of a viable H_0_ hypothesis.

#### 2.4.2. Calculation and usage of temporal signal-to-noise ratio

To gauge the potential impact of nuisance regression on data metrics on a group level we analyzed the development of temporal signal-to-noise ratio (tSNR) as a consequence of noise regression using either RETROICOR, HR, RVT, aCompCor, and AROMA. These calculations were made with a custom Python (v3.11.3) script utilizing the functionality of Nilearn and NiBabel (https://github.com/nipy/nibabel; DOI: 10.5281/zonodo.4295521). Prior to tSNR calculation, images were smoothed using a 6mm Gaussian FWHM-kernel, this was done to align with ICA-AROMA processing, which was run on 6mm smoothed data as originally described by (Pruim, Mennes, van Rooij, et al., 2015).

TSNR was calculated using the formula

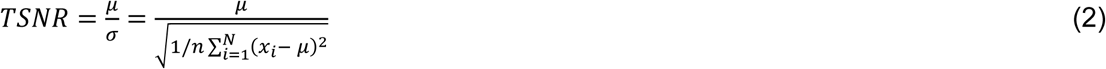

where N is the number of time points in a voxel time-series, µ is the mean of the time series and σ is its standard deviation. Calculation of voxel-wise tSNR was done for the entire functional data matrix, resulting in a tSNR matrix for the functional run. To test tSNR improvement as a result of noise regression, we used the residuals of the previously described GLMs with the respective cleaning procedures for the tSNR calculation. To allow for improved relative comparability, we computed the within-subject increase of tSNR as a consequence of noise correction over no correction in percent. This approach resulted in percent change TSNR maps for each method with RETROICOR, HR, RVT, AROMA, and aCompCor noise regression respectively

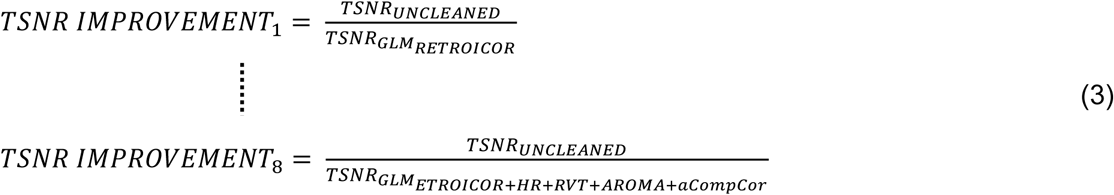

To calculate the unique improvement of tSNR as a consequence of RETROICOR, HR and RVT when combined with AROMA and aCompCor, we calculated a voxel-wise difference score in tSNR as described in Formula 3.

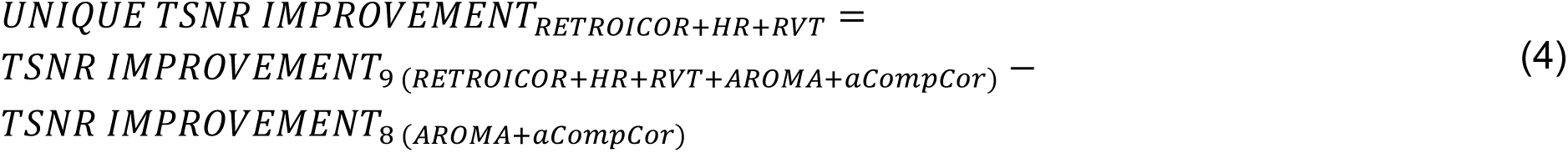

Inferential testing of TSNR improvement was done by means of one-sample t-tests, with FDR-correction for multiple comparisons. These comparisons were made separately for four different regions of interest to account for a different level of abstractions and proximities to physiological noise sources: (1) whole brain, (2) cortical grey matter, (3) brainstem and (4) locus coeruleus.

### 2.5. Identifying potential misclassifications in ICA-AROMA

One aspect of possible individual variation in physiological nuisance regression using ICA-AROMA is the potential of misclassification of physiological noise components as signal in the ICA-AROMA algorithm following ICA-MELODIC. To control whether effects of RETROICOR are caused by component misclassifications in ICA-AROMA we deployed a comparison assessing whether spatial similarity in unique RETROICOR effects as implemented in GLM _RETROICOR+AROMA_ and melodic components classified as a signal, might increase AROMA classification accuracy. RETROICOR was chosen to be used separately here, as it produces a distinct spatial pattern around the brainstem, ventricles and Circle of Willis.

We ran regression models for each MELODIC component on a per-subject basis to create a spatial back projection of a component. Subsequently, a ‘Goodness of Fit’ score was calculated between each melodic component back projection and a masked binarized version of all suprathreshold voxels after FWE-correction uniquely explaining variance across all RETROICOR regressors in GLM_9_. The score was calculated, by comparing the ratio of variance explained by the melodic component inside versus outside the RETROICORs unique mask. All components across subjects that fulfilled the conditions (1) goodness of fit higher than 0.75 and (2) a component being classified as signal, were then taken for further analysis.

Following overlap identification, each identified MELODIC component was added to a subjects’ respective RETROICOR and AROMA regression model (GLM_AROMA+RETROICOR_) to assess whether addition of these potential misclassifications reduces the variance explained by RETROICOR. To quantify this reduction of RETROICORs explanatory power, we calculated a difference value between average z-scores across (after whole-brain FWE-correction) before and after addition of the possibly misclassified component for each subject. This created a mean z-difference score for each possible misclassification.

To get insight into the question whether RETROICORs observed unique explanatory power is dependent on misclassifications, we calculated the correlation between the number of FWE-corrected suprathreshold voxels within the unique RETROICOR maps and the sum of z-difference values per subject when adding potential misclassifications to the model. A possible negative correlation here would hint at a stronger overall reduction in RETROICORs explanatory power the higher the number of ICA-AROMA misclassifications.

### 2.6 Data and code availability

All scripts used for analysis and visualization in this article are available on https://github.com/MKrentz/Methods-for-Physiological-Noise-Correction. Pseudonymized data can be accessed on request in BIDS format from the Donders Repository at https://data.donders.ru.nl. Due to privacy or ethical restrictions, raw MR images are not publicly available.

## 3. Results

### 3.1. tSNR changes and differences resulting from noise regression

To assess the impact of RETROICOR, HR and RVT in comparison to the data-driven correction methods we first visually assessed the spatial distribution of tSNR changes as a consequence of the respective noise removal application. These images allow us to gauge the spatial layout of observed tSNR changes resulting from nuisance regression without the presence of other cleaning methods. Furthermore, improvement is measured in percent and scales are variable for purposes of spatial pattern visualization and not directly comparable between methods.

#### 3.1.1. Spatial layout of tSNR improvements

The spatial distribution of tSNR improvement, as shown in Figure 1, is a purely descriptive depiction of the spatial features of tSNR changes as a consequence of nuisance regression. It shows a typical pattern of RETROICOR related improvement (Figure 1a), with clustering around arterial vascular areas. As expected, strong changes are visible along the basilar artery and cerebral arteries as well as ventricles. Comparatively, the tSNR changes as a consequence of HR (Figure 1b) and RVT (Figure 1c) nuisance regression show low spatial specificity. Peak changes for both a restricted to the edge of the brain, with an occipital peak forming around the sagittal sinus. However, in line with previous findings they show a consistent pattern of grey matter improvement (Chang et al., 2009; Shmueli et al., 2007). aCompCor shows a widespread pattern of tSNR improvement (Figure 1d), with however strong similarity to RETROICORs spatial specificity. While tSNR improvement is overall more homogenous subcortical peaks mimic RETROICOR tSNR improvement by being centered around the basilar artery, Circle of Willis and ventricular system, in line with its origin in CSF principal component extraction. Comparatively, tSNR change as a consequence of AROMA cleaning as visible in Figure 1e show a much more complex spatial distribution. While generally showing much higher absolute tSNR change, in line with its intended use for a variety of noise sources, high tSNR changes can be seen in prefrontal and occipital cortex areas typically associated with head motion (Pruim, Mennes, Buitelaar, et al., 2015; Pruim, Mennes, van Rooij, et al., 2015). However, substantial tSNR changes also occur in most inferior brainstem areas which are strongly affected by inflow artifacts and coil insensitivity and are consequently most likely filtered out via the frequency-based component selection in ICA-AROMA (Pruim, Mennes, van Rooij, et al., 2015). It is apparent that despite its intentionality of motion correction AROMA also strongly improved tSNR in areas associated with RETROICOR.

**Figure 1.**
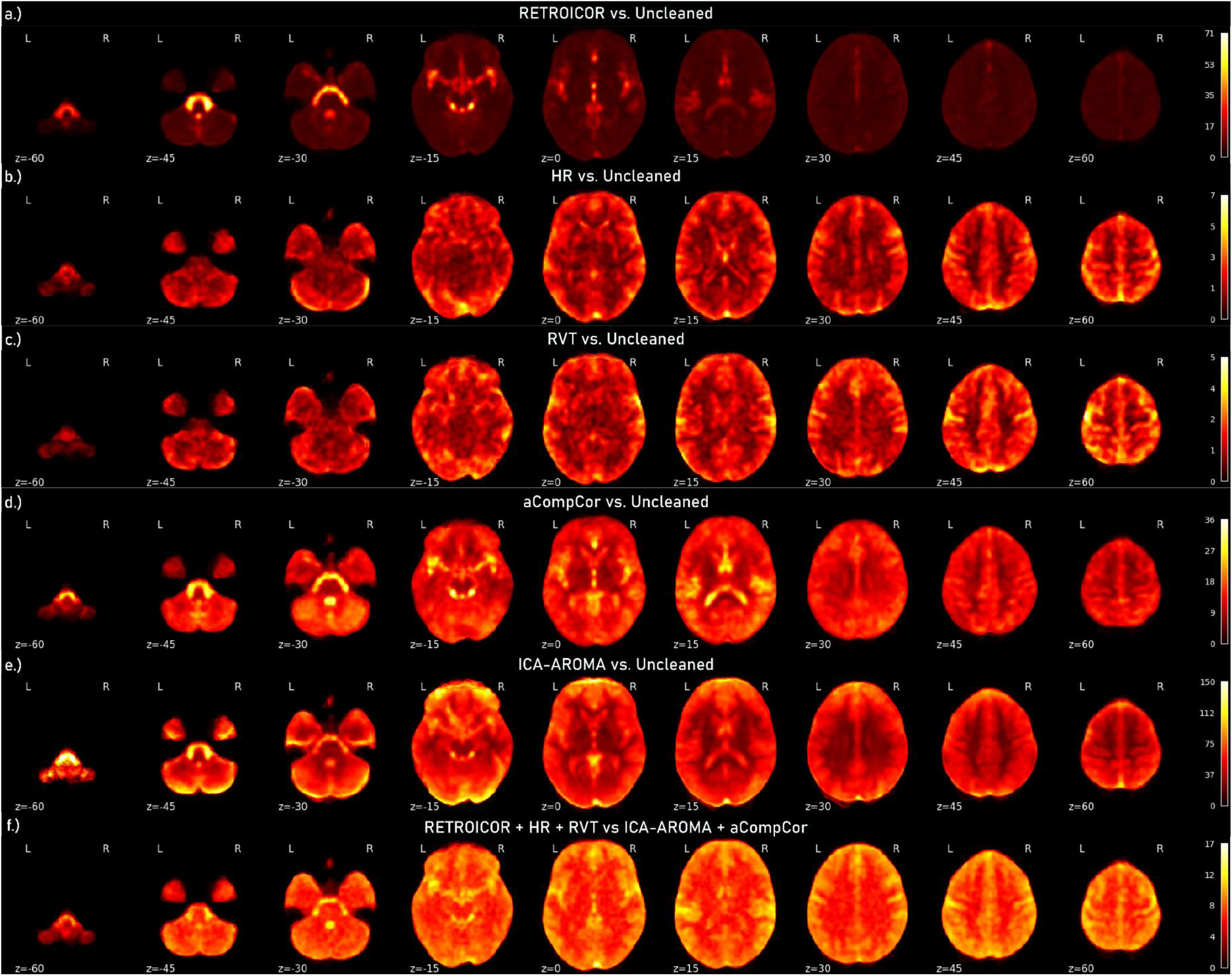
Mean tSNR maps with variable scaling to show spatial pattern of relative improvement of tSNR compared to uncleaned data for all methods individually (a.-e.), as well as the main comparison (f.). In section e.) the threshold was capped at 150, to allow for pattern visibility. Inferior slices exhibited otherwise extreme improvement due to correction of susceptibility effects.

When visualizing combinatory tSNR improvement of the physiological recordings-based techniques in presence of AROMA and aCompCor (Figure 1f), overall tSNR improvement is low yet clear structural specificity around the vascular and ventricular areas hallmark to RETROICOR are visible. Despite a strong visible contribution of both aCompCor and AROMA to tSNR improvement in these areas as visible in Figure 1d and Figure 1e., a unique effect seems to be retained when applying RETROICOR, HR and RVT. Note that no thresholding was implemented for the creation of these images.

#### 3.1.2. Group effects within ROIs

To test the observed group effects on tSNR and the variety in the spatial layout of tSNR changes, we compared the tSNR change based on four different ROIs. These regions were chosen to be progressively smaller and towards the brainstem in higher proximity to areas classically affected by physiological noise. We assessed tSNR change for different masks: whole-brain (1), grey matter (2), brainstem (3) and the locus coeruleus close to the 4^th^ ventricle (4) as visible in Figure 2.

**Figure 2.**
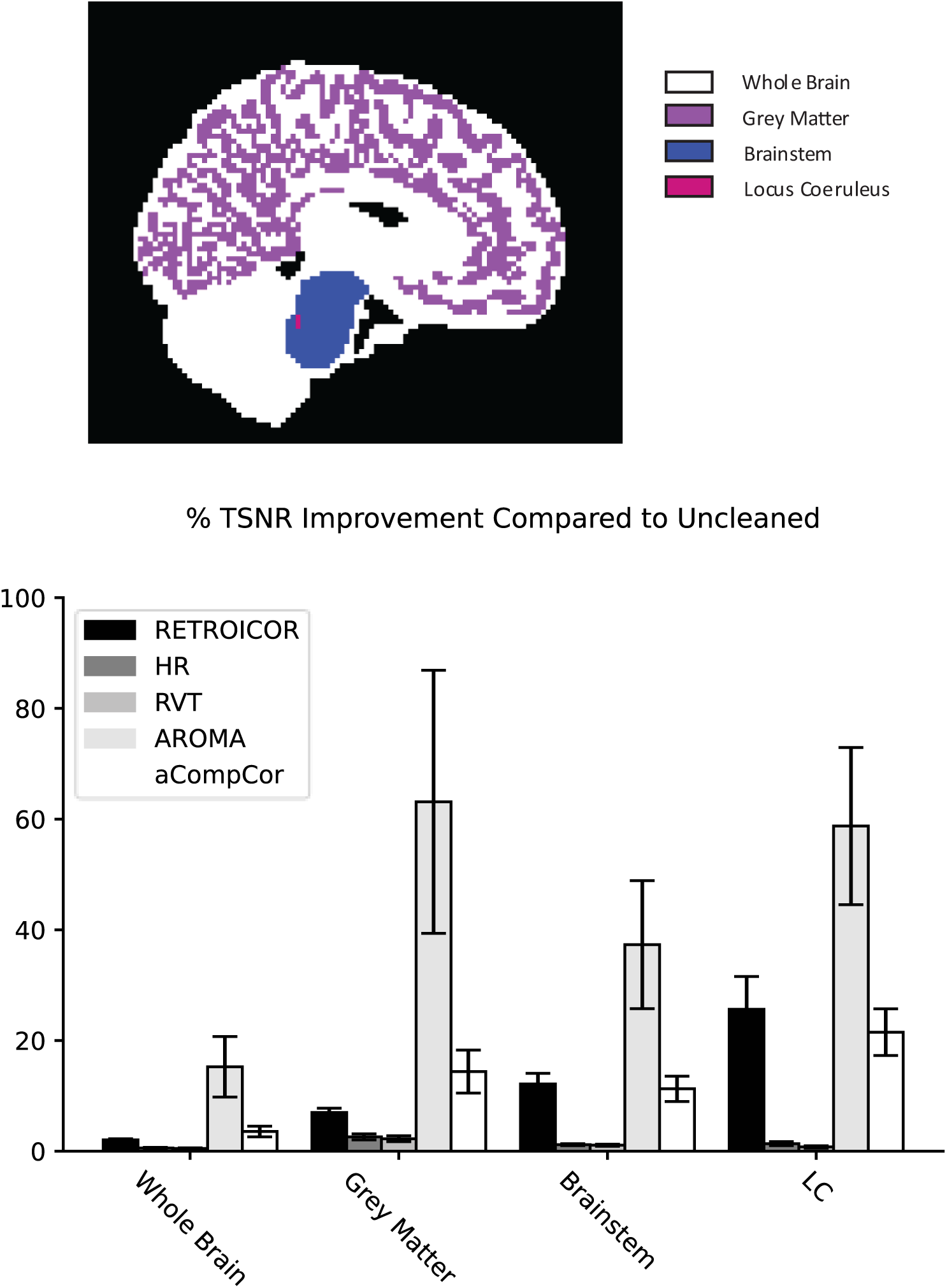
TSNR improvement as a consequence of nuisance regression of RETROICOR, HR, RVT, AROMA, and aCompCor in percent. Error bars represent 95% confidence intervals.

First, as a sanity check, we tested whether the different noise correction methods had a significant impact on tSNR across the four different mask levels as visualized in Figure 2. AROMA shows a consistent strong improvement of tSNR in line with the multi-facetted approach it takes. Similarly, aCompCor and RETROICOR show significant difference from 0 across all ROIs. HR and RVT show a significant reduction in tSNR, yet with absolute lower percentage than the other methods. Respective comparisons between methods were not computed here, as (1) samples across ROIs are not independent and partially redundant and (2) the singular use of one method and its impact on tSNR is of little overall value in respect to the questions of this study. TSNR improvement was consistently impactful across all regions of interest. With RETROICOR increasing tSNR significantly for whole brain (1); *t*(25) = 19.11, *p* < .001, *d* = 3.748; cortical grey matter (2); *t*(25) = 17.433, *p* < .001, *d* = 3.419; brainstem (3); *t*(25) = 12.0786, *p* < .001, *d* = 2.369; as well as locus coeruleus (4); *t*(25) = 8.338, *p* < .001, *d* = 1.635. HR regression significantly increased tSNR with the regions of interest with whole brain (1); *t*(25) =10.714, *p* < .001, *d* = 2.101; cortical grey matter (2); *t*(25) = 9.069, *p* < .001, *d* = 1.779; brainstem (3); *t*(25) = 13.465, *p* < .001, *d* = 2.641; and locus coeruleus (4); *t*(25) = 7.169, *p* < .001, *d* = 1.406. Similarly, RVT significantly increased tSNR for whole brain (1); *t*(25) = 9.202, *p* < .001, *d* = 1.805; cortical grey matter (2); *t*(25) = 8.053, *p* < .001, *d* = 1.579; brainstem (3); *t*(25) = 11.385, *p* < .001, *d* = 2.233; as well as locus coeruleus (4); *t*(25) = 6.499, *p* < .001, *d* = 1.275. AROMA regression led to significant tSNR improvement across regions of interest for whole brain (1); *t*(25) = 5.355, *p* < .001, *d* = 1.050; cortical grey matter (2); *t*(25) = 5.103, *p* < .001, *d* = 1; brainstem (3); *t*(25) = 6.2, *p* < .001, *d* = 1.216; and locus coeruleus (4); *t*(25) = 7.953, *p* < .001, *d* = 1.56. Furthermore, aCompCor does significantly explain variance for whole brain (1); *t*(25) = 7.083, *p* < .001, *d* = 1.389; cortical grey matter (2); *t*(25) = 7.113, *p* < .001, *d* = 1.395; as well as brainstem; *t*(25) = 9.4719, *p* < .001, *d* = 1.858; and locus coeruleus (4); *t*(25) = 9.812, *p* < .001. *d* = 1.924.

Figure 3 displays the tSNR improvement unique to an addition of RETROICOR, HR and RVT to a combination of both data-driven methods. Testing the difference shows that the physiological data derived methods uniquely improves tSNR over AROMA and aCompCor, showing effects across all ROIs for whole brain (1); *t*(25) = 44.961, *p* < .001, *d* = 8.818; cortical gray matter; *t*(25) = 39.189, *p* < .001, *d* = 7.685; brainstem; *t*(25) = 55.499, *p* < .001, *d* = 10.884; and locus coeruleus; *t*(25) = 13.099, *p* < .001, *d* = 2.569.

**Figure 3.**
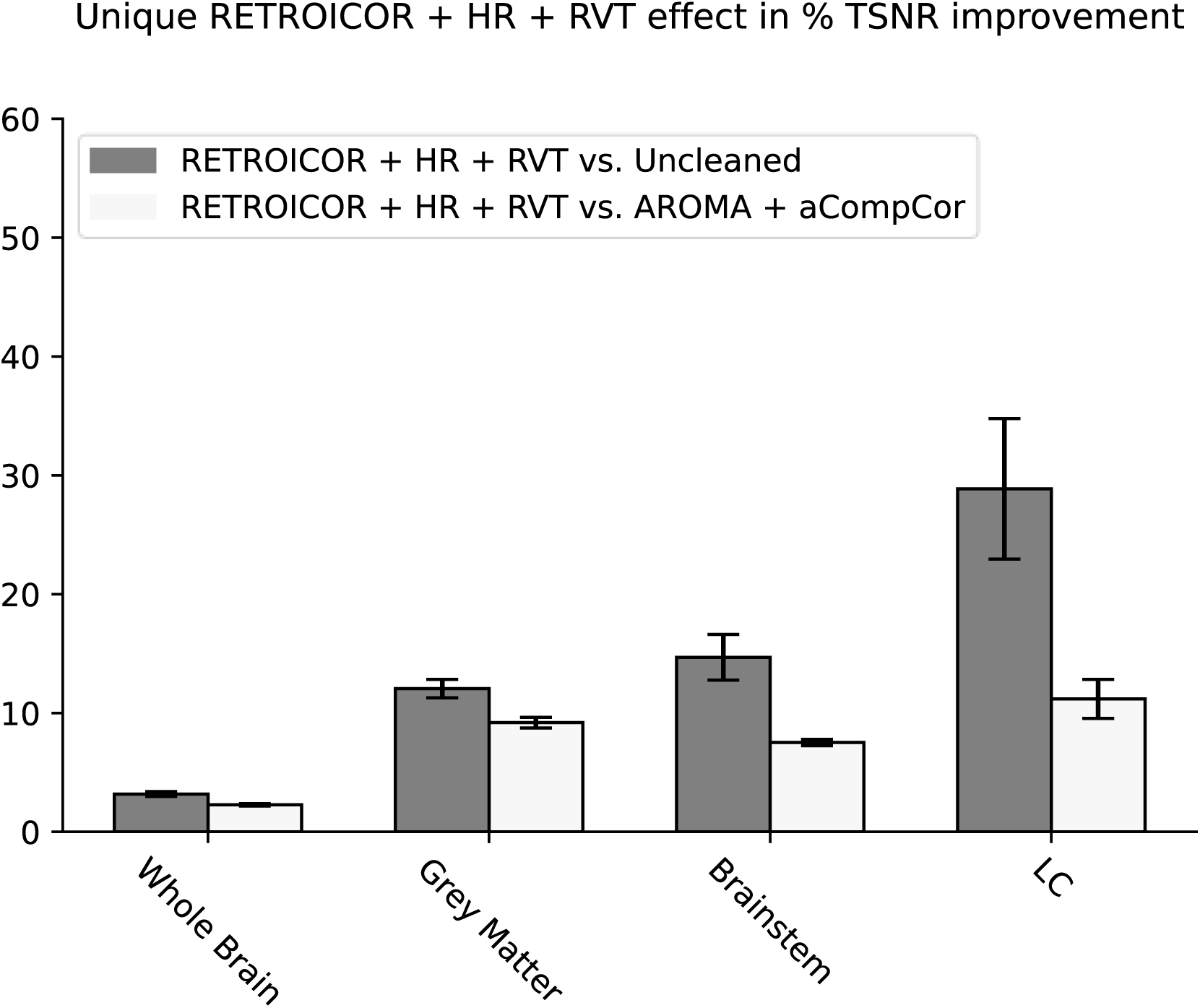
Percent TSNR improvement of RETROICOR, HR and RVT nuisance regression compared to the unique TSNR improvement of RETROICOR, HR and RVT when tested versus AROMA and aCompCor in Percent. Error bars represent 95% confidence intervals.

### 3.2. Statistical improvement for selection of noise regression methods

To directly assess the inter-individual consistency of the different noise correction procedures, the computed GLMs were chosen to test the variance explained by the different noise correction methods in a resting state run for RETROICOR, HR, RVT, ICA-AROMA, and aCompCor respectively. The calculations were made for all methods separately in comparison to uncleaned data (Appendix C1-C5). Spatial patterns are generally consistent with tSNR maps as seen in Figure 1 yet generally differ in the amount of variance explained across methods as well as inter-subject consistency.

#### 3.2.1. tSNR improvement due to RETROICOR, HR and RVT over ICA-AROMA and aCompCor

Investigating the unique variance of combined RETROICOR, HR and RVT when testing with a combination of AROMA and aCompCor a pattern appears showing high inter-subject variance in the impact of physiological recording-based nuisance regression, as can be seen in Figure 4. As visible on subjects 006, 012, 015, 024 and 025, AROMA in combination with aCompCor does well in explaining physiological noise. However, in contrast, more impacted subjects (002, 003, 007, 008, 016, 017, 023, 027, 029, 030), retain a significant amount of uniquely explained variance by RETROICOR, HR and RVT. While the impact of RETROICOR, HR and RVT on an individual level is comparatively decreased, especially around the brainstem as well as third and fourth ventricles, the combination of data-driven approaches does not appear to consistently account for all physiological noise captured with peripheral recordings.

**Figure 4.**
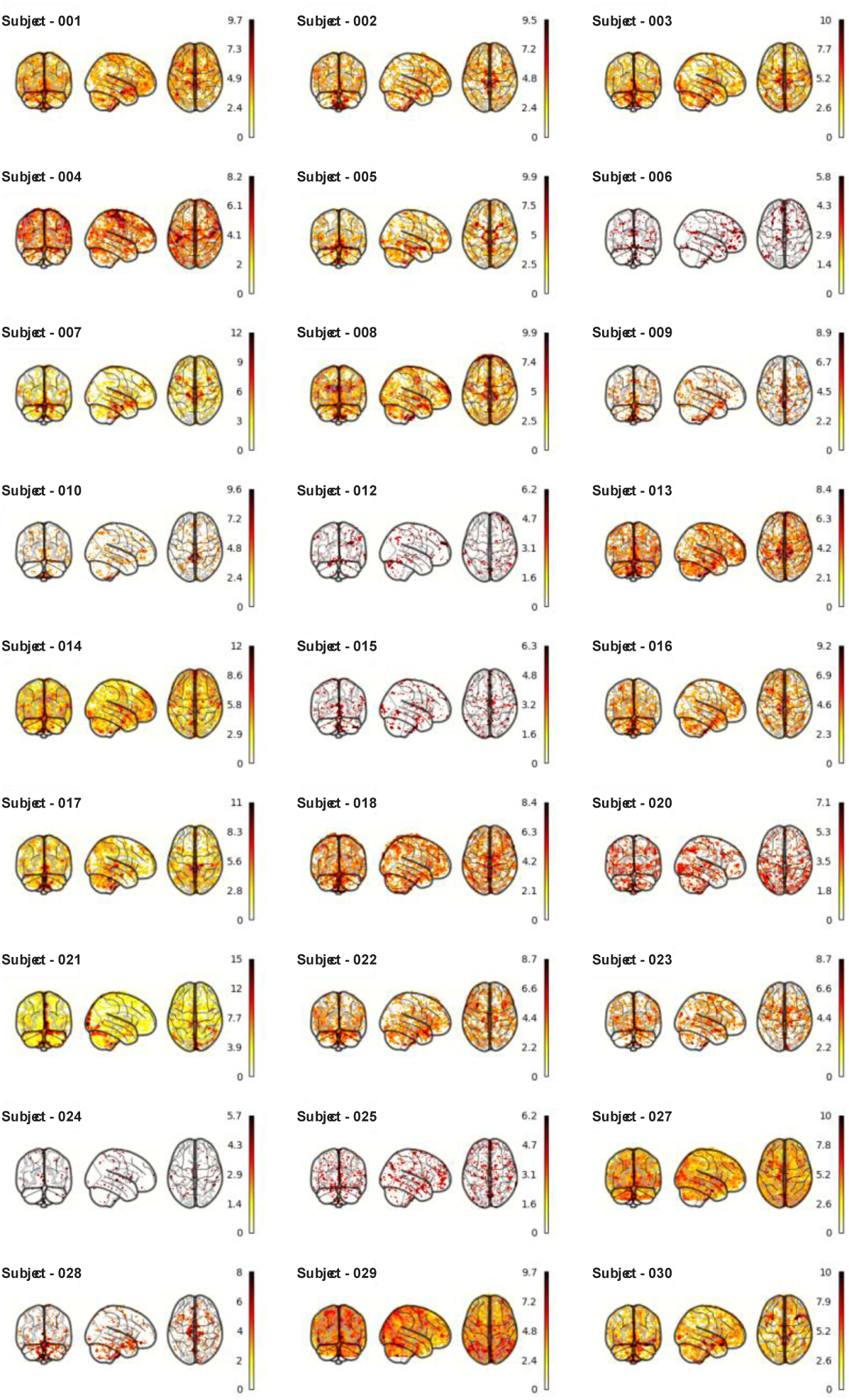
Subject-wise spatial representation of individually explained variance uniquely attributed to RETROICOR, HR and RVT with AROMA and aCompCor in the model (thresholded at pFDR <.05).

### 3.3. Misclassifications in ICA-AROMA

One reason for inconsistency of data-driven methods could be the classification of noise components as signal using a pre-specified set of rules for noise identification in ICA-AROMA. It is possible that variability in the consistency of this classification process, due to it not being tailored for physiological noise specifically, accounts for some of the physiological recording-based effects. Consequently, we tested whether the spatial maps of our RETROICOR approach correspond to spatial maps of ICA components classified as signal by ICA-AROMA. Figure 5 shows an example of a participant for whom the addition of a component initially misclassified as signal severely reduces the variance explained by RETROICOR.

**Figure 5.**
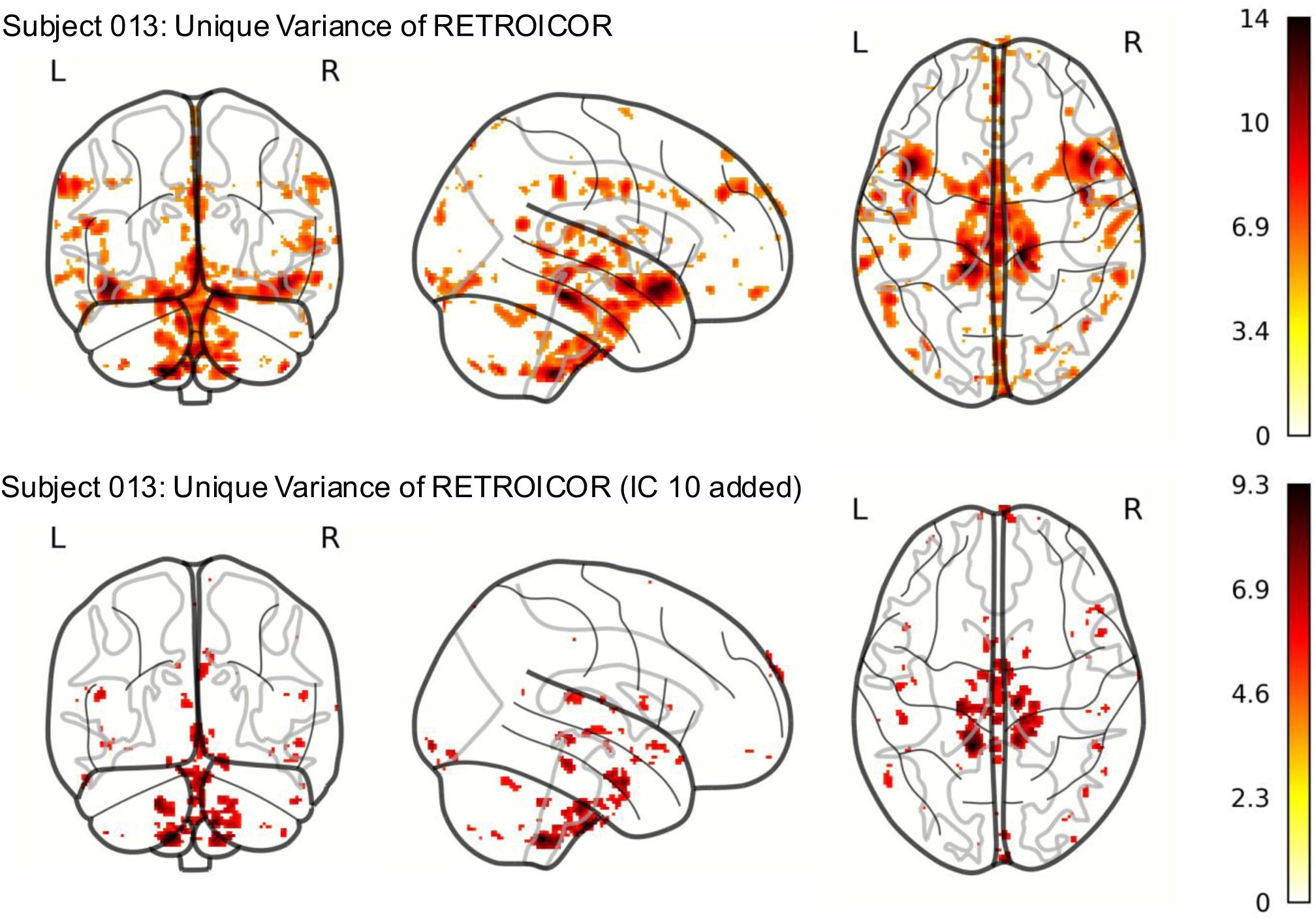
Unique explained variance of RETROICOR after adding an IC initially misclassified as signal to the array of AROMA regressors for subject 013.

As shown in Figure 6, no significant Spearman rank-order correlation was observed between the individual sum of z-differences (resulting from adding all possible misclassifications to a subject’s RETROICOR model) and the magnitude of the unique RETROICOR effect, *ρ* = -0.15, *p* = .49. This suggests that inconsistencies in ICA-AROMA component classification are not the sole driver of RETROICOR’s ability to explain additional variance beyond ICA-AROMA.

**Figure 6.**
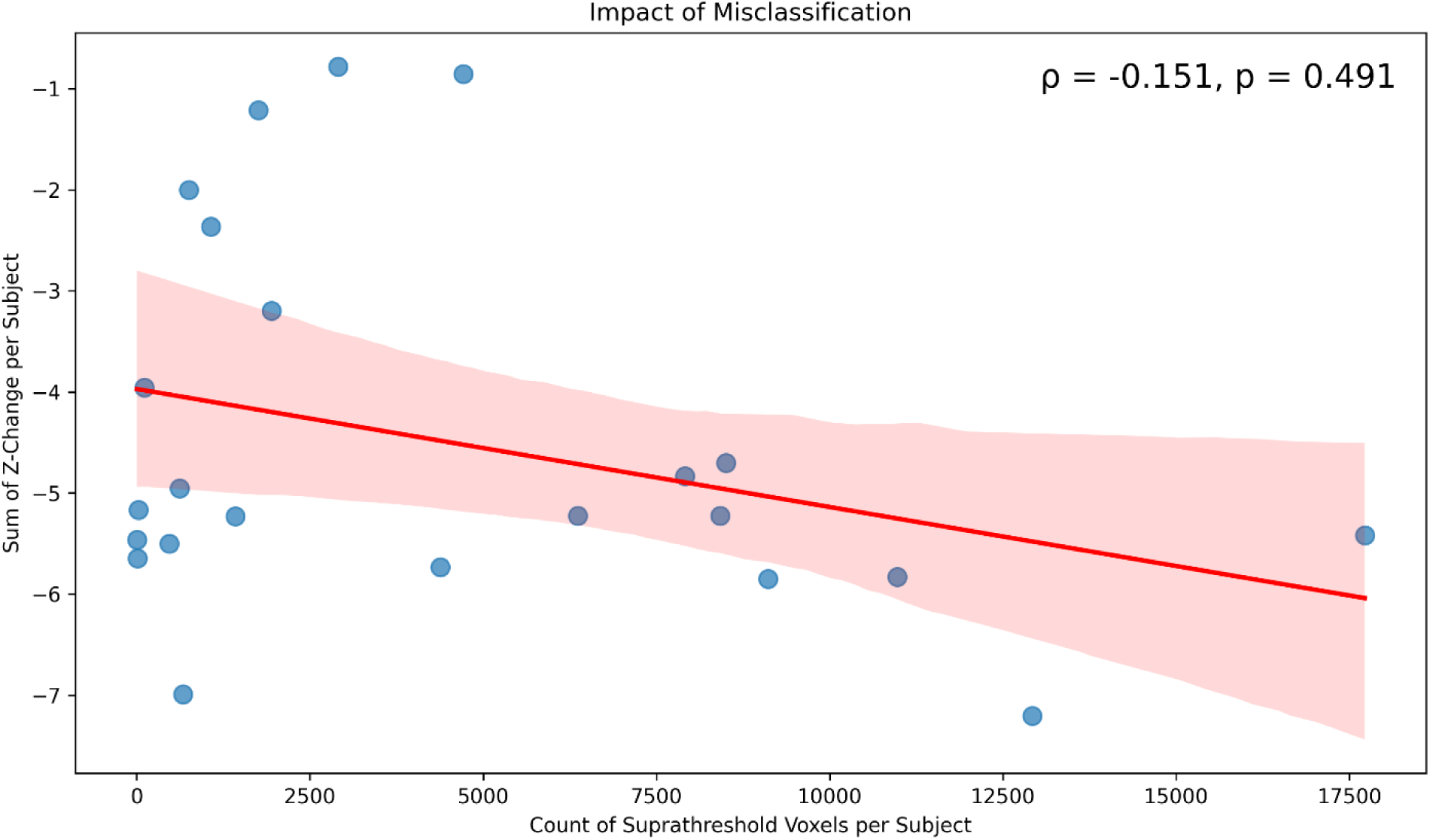
Association between sum of overall z-change for all misclassifications per subject and the total number of FWE-corrected suprathreshold voxels within contrast explaining unique variance of RETROICOR over ICA-AROMA. Error band represents 95% confidence intervals.

Consequently, correcting misclassifications by reclassifying MELODIC components as noise does not fully account for RETROICOR’s retained explanatory power. A further limitation arises from the visual identification of components with high “Goodness of Fit” and strong z-changes that nonetheless contain meaningful signal. These components reduce RETROICOR’s effects, implying they may encode redundant physiological noise not captured by ICA-AROMA.

## 4. Discussion

The need and acceptance for streamlined pre-processing pipelines has been growing. The now commonly used pre-processing tool fMRIPrep can automatically output regressors to be included in GLM analysis of functional MRI data with aCompCor and allow for easily accessible computation of ICA-AROMA to remove noise attributed to a variety of sources (Esteban, Markiewicz, et al., 2018). This approach is compelling in comparison to classical physiological noise correction methods utilizing peripheral measurements, such as RETROICOR, HR or RVT (Glover et al., 2000). Peripheral physiological measures require additional processing steps, equipment, and preparation during the actual execution of an experiment. The question arises how applicable the combination of data driven methods is to account for physiological noise in noise sensitive settings such as brainstem imaging using high-resolution imaging. As expected, all tested methods (i.e. RETROICOR, HR, RVT, ICA-AROMA and aCompCor) show a significant impact on tSNR as well as explained variance (Behzadi et al., 2007; Krüger & Glover, 2001; Pruim, Mennes, van Rooij, et al., 2015). While aCompCor and ICA-AROMA show a generalized high amount of explained variance, RETROICOR shows a consistent impact on explained variance, especially around fluid cavities such as ventricles and arteries surrounding the brainstem. HR and RVT show lower overall changes as well as less spatial specificity, being more widespread across mostly grey matter. On a group level, the difference between the data-derived methods and physiological recording methods appears small, yet a look at individual expressions reveals a consistent impact of peripherally recorded physiology with some inconsistencies in ICA-AROMA’s and aCompCor’s combined predictive ability.

### 4.1. Effects of noise nuisance regression on group level

All tested methods show unique improvements in tSNR in comparison to a baseline that differs both in magnitude as well as spatial profile. In line with prior findings around RETROICOR, a large part of the removed variance can be attributed to both the cardiac and the respiratory cycle. These tSNR improvements are largely explained by the underlying removal of non-BOLD physiological noise arising from cardiac pulsations. These motion-like pulsations result in changes mostly visible in fluid-filled cavities such as the 4^th^ and 3^rd^ ventricles, as well as arteries and the Circle of Willis (Glover et al., 2000; Krüger & Glover, 2001). In addition to RETROICOR, two other methods based on peripheral physiological data, HR and RVT, showed significant contributions to tSNR improvement with less spatial specificity across grey matter and along brain edges combating cycle-independent low-frequency physiological noise-related changes. In contrast, ICA-AROMA reveals a more widespread improvement of tSNR at a higher magnitude in line with its broader scope of removal of both physiological and motion noise (Pruim, Mennes, Buitelaar, et al., 2015; Pruim, Mennes, van Rooij, et al., 2015). While less specific to pulsatility-related noise as RETROICOR, it yet shows high improvement of tSNR in areas also impacted by RETROICOR cleaning. Peak improvement is centered on most inferior slices, likely heavily impacted by high-frequency noise due to both coil insensitivity as well as susceptibility artifacts, which thus is likely captured by ICA-AROMAs high-frequency content selection criteria (Pruim, Mennes, van Rooij, et al., 2015). Similarly, it is dominated in prefrontal and occipital locations related to bulk head movement as well as susceptibility artifacts (Pruim, Mennes, Buitelaar, et al., 2015). aCompCor in comparison shows a similar focus on CSF and cardiac areas as RETROICOR (Muschelli et al., 2014). The PCA-based approach shows overall less specificity to high pulsation areas yet nonetheless resembles in its peaks the previously shown pattern of RETROICOR (i.e. regions surrounding 3^rd^ and 4^th^ ventricle as well as Circle of Willis). The overall improvement of tSNR centered around gray matter is furthermore an expected consequence of the relatively broad approach of white matter nuisance regression (Muschelli et al., 2014). While the TR of 2.02s leads to aliasing of actual cardiac frequency, it appears as if spatial coherence of the PCA components can compensate to some degree achieving removal of physiological noise even with relatively long TR (T. T. Liu, 2016). If aliasing would have a strong impact, it should affect cardiac pulsations over respiration-induced pulsatility (Huotari et al., 2019), yet the relatively long TR in this study does not negate the general detectability of pulsation-related signal changes.

When adding ICA-AROMA and aCompCor to a GLM, the unique tSNR improvement of the peripheral recording-based methods is generally strongly reduced. However, while large parts of its value are captured by these data-derived methods, small improvements remain, specifically centered around aforementioned fluid cavities affected by high cardiac pulsatility as well as occipital effects related to venous drainage. As expected, RETROICOR, HR and RVT retain a high degree of improvement closer to areas of pulsation, such as the LC, directly at the edge of the 4^th^ ventricle. In addition, they demonstrate notable improvements not only near areas of pulsation but also throughout grey matter and the entire brainstem. As mechanistically there is reason to expect a difference in effectiveness and an upside to direct nuisance regression over data-derived components in this setting, we investigated whether variation in inter-individual consistency can account for the found differences.

### 4.2. Effects of noise nuisance regression on individual level

For all methods, a pattern consistent with expected spatial features of variance explained was visible when testing in absence of other correction methods (Appendix C1-C5), though some features appeared more consistent than others hinting at variability in individual effectiveness.

This picture of individual variability in the effectiveness of noise removal becomes more apparent when comparing RETROICOR, HR and RVT to ICA-AROMA and aCompCor. For some subjects, ICA-AROMA and aCompCor explain large amounts of variance in areas sensitive to cardiac pulsations, and thus almost entirely negate the effect of physiological recording-based confound removal. However, in other subjects, ICA-AROMA and aCompCor fail to account for the same type of variance uniquely identified by RETROICOR, HR and RVT. Thus, it can be inferred that while effective in removing physiological noise, inter-individual inconsistency creates a problem – one that could be detrimental when trying to assess areas, such as the brainstem, strongly susceptible to cardiac and respiration-induced pulsations. There are multiple possible explanations for these individual inconsistencies.

One potential source of inconsistency in ICA-AROMA is the misclassification of components, particularly when physiological noise is incorrectly identified as signal. Although ICA-AROMA was not specifically designed to target physiological noise, both our findings and those of others indicate that it holds promise as an effective physiological denoising tool. However, as demonstrated here, misclassification of ICA components can substantially impact the interpretability of resting-state fMRI data. Notably, the absence of a negative correlation between the unique variance explained by RETROICOR after correcting for misclassifications and its overall explanatory power suggests that RETROICOR and ICA-AROMA may target similar physiological noise components, though RETROICOR may model these aspects more precisely. Thus, RETROICOR’s effectiveness is unlikely to be solely a result of ICA-AROMA misclassification but rather reflects its unique capacity to correct for physiological noise.

One alternative is to use methods such as ICA-FIX to train and select noise components with higher accuracy and consistency. However, ICA-AROMA provides a degree of reproducibility when continuously implemented as part of a common pre-processing pipeline and does not require the training of a classifier and accompanying expertise in doing so. Consequently, adjustments could be made, using similar methods as used in this article when identifying misclassifications, to create an additional selection criterion based on overlap with known physiological noise components (Brooks et al., 2013; T. T. Liu, 2016).

Another issue of inconsistency could be related to the nature of the sequence used in this study. Brainstem imaging requires compromises to achieve relevant resolution and consistency. As the original focus of this study was the imaging of brainstem nuclei under externally perturbated manipulation, the used TR of 2.02 seconds with a relatively short experimental runtime – a trade-off to achieve usable spatial resolution and quality – could hurt the formation of consistent strong ICA components (Pruim, Mennes, Buitelaar, et al., 2015; Van Schuerbeek et al., 2022). As ICA component numbers rise as a consequence of increasing volume numbers, it has been shown that smaller functional brain networks only separate into unique components under ideal circumstances (Cohen et al., 2021). Therefore, it could be the case here that aspects of physiological noise are inconsistently included in large components comprising mostly signals of interest.

Similar problems arise for aCompCor in this setting as well as the chosen mode of component selection. The chosen methods of including the first six principal components created by aCompCor are detached from the explanatory power of these components and do not account for the actual total amount of variance explained. It can thus be that these components have varying explanatory impacts between subjects. This aspect can play an important role in the individual variation of explanatory power. An alternative method, often referred to as aCompCor50, similarly calculated by default in fMRIPrep, includes all components up to 50% of the total explained variance (Behzadi et al., 2007; Muschelli et al., 2014). When under the assumption of white matter and CSF signal as noise this would include the most coherent temporal and spatial constructs, it would also lead to an enormous decrease in tDOF on average. When dealing with a low number of volumes, this could have a further detrimental impact on the ability to draw statistical inference. Other criteria, such as manual cut-off based on leveling plotted explanatory value, does both add increased labor and an undesired increase in manual input, but similarly also relies on TR, as the quality of component formation has been shown to increase with lower TRs and higher volume number (Cohen et al., 2021). While methods have been developed to create a more integrated combination of ICA-AROMA and aCompCor (van Schuerbeek et al., 2022), doubt has similarly been cast on the assumption that white matter signal is completely devoid of grey matter signal of interest (Gawryluk et al., 2014). Based on these limitations, it is clear that ICA-AROMA and aCompCor offer an easy and convenient array of tools to account for different types of noise. However, when tackling regions, such as brainstem nuclei, that are highly sensitive to cardiac pulsations, the need for accuracy and consistency in combination with additional demands on sequence parameters, should justify the addition of peripheral physiology recordings. When choosing higher volume numbers and lower repetition times the specificity might increase, yet at a potential detrimental trade-off for usability in brain-stem imaging.

### 4.3. Limitations of the current study

Even though the strength of peripheral measurements is agnostic to features of the sequence, the actual accuracy of the data-derived methods was not assessed in a competitive setting. This article only looks at the impact of physiological noise in a resting-state setting and thus ignores problems arising from combinations of tasks and nuisance variables. Classically used ICA-AROMA is able to remove noise, yet does so irrespective of task regressors, if not introduced into GLMs as nuisance variables. We did not use variation of a data proxy such as functional connectivity to gauge the impact of physiological noise correction. Thus, for brainstem regions it is unclear how the used correction methods affect the usability of derived data, whether being meaningful signal or dominated by noise. While it does not diminish the results of this study, it emphasizes that the statements here are aiming towards feasibility and not usability. An evaluation of possible interaction between e.g. ICA-AROMAs inconsistency in removal and formation of brain networks is beyond the scope of this article. Furthermore, within this study, we did not assess multiple sessions and thus draw no inference about inter-session stability of RETROICOR, HR, RVT, ICA-AROMA or aCompCor nuisance regression. The conclusions drawn here are thus only applicable to a context that was based on a trade-off scenario for brainstem imaging.

### 4.4. Conclusion

Overall, we show the effectiveness of RETROICOR, HR, RVT, ICA-AROMA, and aCompCor to increase tSNR as a consequence of accounting for noise. While easily accessible methods such as ICA AROMA and aCompCor can account for most of the explanatory value of methods using peripherally recorded physiology, a small subset of unique information remains, partially arising from higher intra-group consistency of noise detection. Apparent instability in ICA-AROMAs ability to detect physiological noise in this study points towards desired additional reliance on peripheral measures, even when combined with tools to tackle physiological noise such as aCompCor. Hence, adding RETROICOR, HR and RVT can increase confidence in noise removal stability and can furthermore impact data quality when assessing brainstem areas or areas in proximity to fluid cavities.

## Acknowledgements

This work was supported by a grant from the European Research Council (ERC-2015-CoG 682591).

## Author Contributions

**Martin Krentz**: Conceptualization, Methodology, Software, Formal analysis, Investigation, Project administration, Writing - Original draft. **Rayyan Tutunji**: Conceptualization, Writing - Review & Editing. **Nikos Kogias**: Investigation, Writing – Review & Editing. **Hariharan Murali Mahadevan**: Project administration, Investigation, Data Curation. **Zala C. Reppmann**: Project administration, Investigation, Data Curation. **Florian Krause**: Conceptualisation, Methodology, Supervision, Writing – Review & Editing. **Erno J. Hermans**: Conceptualisation, Methodology, Supervision, Writing – Review & Editing, and Funding acquisition.

## Notes

### Competing Interest Statement

The authors have declared no competing interest.

### Summary of Updates

Title updated to better reflect manuscript focus; General language updates; Manuscript expanded to include heart rate and respiratory volume per unit time across analysis; Methods and analysis section updated to account for an initial inaccuracy in the computation of RETROICOR regressors; Updated analysis of ICA-AROMA misclassification (Section 2.5 & 3.3); Revised Figures 1-7 as well as C1-C5 based on expanded analysis.

## References

Abraham, A., Pedregosa, F., Eickenberg, M., Gervais, P., Mueller, A., Kossaifi, J., Gramfort, A., Thirion, B., & Varoquaux, G. (2014). Machine learning for neuroimaging with scikit-learn. Frontiers in Neuroinformatics, 8. 10.3389/fninf.2014.00014

Avants, B. B., Epstein, C. L., Grossman, M., & Gee, J. C. (2008). Symmetric diffeomorphic image registration with cross-correlation: Evaluating automated labeling of elderly and neurodegenerative brain. Medical Image Analysis, 12(1), 26–41. 10.1016/j.media.2007.06.004

Beckmann, C. F., DeLuca, M., Devlin, J. T., & Smith, S. M. (2005). Investigations into resting-state connectivity using independent component analysis. Philosophical Transactions of the Royal Society of London. Series B, Biological Sciences, 360(1457), 1001–1013. 10.1098/RSTB.2005.1634

Beckmann, C. F., & Smith, S. M. (2004). Probabilistic Independent Component Analysis for Functional Magnetic Resonance Imaging. IEEE Transactions on Medical Imaging, 23(2), 137–152. 10.1109/TMI.2003.822821

Behzadi, Y., Restom, K., Liau, J., & Liu, T. T. (2007). A component based noise correction method (CompCor) for BOLD and perfusion based fMRI. NeuroImage, 37(1), 90–101. 10.1016/j.neuroimage.2007.04.042

Berntson, G. G., Cacioppo, J. T., & Quigley, K. S. (1993). Respiratory sinus arrhythmia: Autonomic origins, physiological mechanisms, and psychophysiological implications. Psychophysiology, 30(2), 183–196. 10.1111/J.1469-8986.1993.TB01731.X

Birn, R. M., Diamond, J. B., Smith, M. A., & Bandettini, P. A. (2006). Separating respiratory-variation-related fluctuations from neuronal-activity-related fluctuations in fMRI. NeuroImage, 31(4), 1536–1548. 10.1016/J.NEUROIMAGE.2006.02.048

Birn, R. M., Murphy, K., & Bandettini, P. A. (2008). The effect of respiration variations on independent component analysis results of resting state functional connectivity. Human Brain Mapping, 29(7), 740. 10.1002/HBM.20577

Brooks, J. C. W., Faull, O. K., Pattinson, K. T. S., & Jenkinson, M. (2013). Physiological Noise in Brainstem fMRI. Frontiers in Human Neuroscience, 0(OCT), 623. 10.3389/FNHUM.2013.00623

Caballero-Gaudes, C., & Reynolds, R. C. (2017). Methods for cleaning the BOLD fMRI signal. NeuroImage, 154, 128–149. 10.1016/J.NEUROIMAGE.2016.12.018

Chang, C., Cunningham, J. P., & Glover, G. H. (2009). Influence of heart rate on the BOLD signal: The cardiac response function. NeuroImage, 44(3), 857–869. 10.1016/J.NEUROIMAGE.2008.09.029

Chu, W. T., Mitchell, T., Foote, K. D., Coombes, S. A., & Vaillancourt, D. E. (2021). Functional imaging of the brainstem during visually-guided motor control reveals visuomotor regions in the pons and midbrain. NeuroImage, 226, 117627. 10.1016/J.NEUROIMAGE.2020.117627

Ciric, R., Thompson, W. H., Lorenz, R., Goncalves, M., MacNicol, E., Markiewicz, C. J., Halchenko, Y. O., Ghosh, S. S., Gorgolewski, K. J., Poldrack, R. A., & Esteban, O. (2022). TemplateFlow: FAIR-sharing of multi-scale, multi-species brain models. Nature Methods, 19, 1568–1571. 10.1038/s41592-022-01681-2

Cohen, A. D., Yang, B., Fernandez, B., Banerjee, S., & Wang, Y. (2021). Improved resting state functional connectivity sensitivity and reproducibility using a multiband multi-echo acquisition. NeuroImage, 225, 117461. 10.1016/J.NEUROIMAGE.2020.117461

Colizoli, O., de Gee, J. W., van der Zwaag, W., & Donner, T. H. (2022). Functional magnetic resonance imaging responses during perceptual decision-making at 3 and 7 T in human cortex, striatum, and brainstem. Human Brain Mapping, 43(4), 1265–1279. 10.1002/HBM.25719

Dagli, M. S., Ingeholm, J. E., & Haxby, J. V. (1999). Localization of cardiac-induced signal change in fMRI. NeuroImage, 9(4), 407–415. 10.1006/NIMG.1998.0424

Dale, A. M., Fischl, B., & Sereno, M. I. (1999). Cortical Surface-Based Analysis: I. Segmentation and Surface Reconstruction. NeuroImage, 9(2), 179–194. 10.1006/nimg.1998.0395

Das, A., Murphy, K., & Drew, P. J. (2021). Rude mechanicals in brain haemodynamics: non-neural actors that influence blood flow. Philosophical Transactions of the Royal Society B: Biological Sciences, 376(1815), 20190635. 10.1098/RSTB.2019.0635

Dreha-Kulaczewski, S., Joseph, A. A., Merboldt, K. D., Ludwig, H. C., Gärtner, J., & Frahm, J. (2015). Inspiration Is the Major Regulator of Human CSF Flow. Journal of Neuroscience, 35(6), 2485–2491. 10.1523/JNEUROSCI.3246-14.2015

Elman, J. A., Panizzon, M. S., Hagler, D. J., Jr., Eyler, L. T., Granholm, E. L., Fennema-Notestine, C., Lyons, M. J., McEvoy, L. K., Franz, C. E., Dale, A. M., & Kremen, W. S. (2017). Task-evoked pupil dilation and BOLD variance as indicators of locus coeruleus dysfunction. Cortex; a Journal Devoted to the Study of the Nervous System and Behavior, 97, 60. 10.1016/J.CORTEX.2017.09.025

Esteban, O., Blair, R., Markiewicz, C. J., Berleant, S. L., Moodie, C., Ma, F., Isik, A. I., Erramuzpe, A., Kent, J. D., Goncalves, M., DuPre, E., Sitek, K. R., Gomez, D. E. P., Lurie, D. J., Ye, Z., Poldrack, R. A., & Gorgolewski, K. J. (2018). fMRIPrep 23.0.2. Software. 10.5281/zenodo.852659

Esteban, O., Markiewicz, C., Blair, R. W., Moodie, C., Isik, A. I., Erramuzpe Aliaga, A., Kent, J., Goncalves, M., DuPre, E., Snyder, M., Oya, H., Ghosh, S., Wright, J., Durnez, J., Poldrack, R., & Gorgolewski, K. (2019). fMRIPrep: a robust preprocessing pipeline for functional MRI. Nature Methods, 16, 111–116. 10.1038/s41592-018-0235-4

Esteban, O., Markiewicz, C. J., Blair, R. W., Moodie, C. A., Isik, A. I., Erramuzpe, A., Kent, J. D., Goncalves, M., DuPre, E., Snyder, M., Oya, H., Ghosh, S. S., Wright, J., Durnez, J., Poldrack, R. A., & Gorgolewski, K. J. (2018). fMRIPrep: a robust preprocessing pipeline for functional MRI. Nature Methods 2018 16:1, 16(1), 111–116. 10.1038/s41592-018-0235-4

Evans, A. C., Janke, A. L., Collins, D. L., & Baillet, S. (2012). Brain templates and atlases. NeuroImage, 62(2), 911–922. 10.1016/j.neuroimage.2012.01.024

Fonov, V. S., Evans, A. C., McKinstry, R. C., Almli, C. R., & Collins, D. L. (2009). Unbiased nonlinear average age-appropriate brain templates from birth to adulthood. NeuroImage, 47, *Supplement* *1*, S102. 10.1016/S1053-8119(09)70884-5

Gawryluk, J. R., Mazerolle, E. L., & D’Arcy, R. C. N. (2014). Does functional MRI detect activation in white matter? A review of emerging evidence, issues, and future directions. Frontiers in Neuroscience, *0*(8 JUL), 239. 10.3389/FNINS.2014.00239/ABSTRACT

German, D. C., Walker, B. S., Manaye, K., Smith, W. K., Woodward, D. J., & North, A. J. (1988). The human locus coeruleus: Computer reconstruction of cellular distribution. Journal of Neuroscience, 8(5), 1776–1788. 10.1523/jneurosci.08-05-01776.1988

Glover, G. H., Li, T.-Q., & Ress, D. (2000). Image-Based Method for Retrospective Correction of Physiological Motion Effects in fMRI: RETROICOR. 10.1002/1522-2594

Gorgolewski, K. J., Burns, C. D., Madison, C., Clark, D., Halchenko, Y. O., Waskom, M. L., & Ghosh, S. S. (2011). Nipype: A flexible, lightweight and extensible neuroimaging data processing framework in Python. Frontiers in Neuroinformatics, 5, 13. 10.3389/FNINF.2011.00013/ABSTRACT

Gorgolewski, K. J., Esteban, O., Markiewicz, C. J., Ziegler, E., Ellis, D. G., Notter, M. P., Jarecka, D., Johnson, H., Burns, C., Manhães-Savio, A., Hamalainen, C., Yvernault, B., Salo, T., Jordan, K., Goncalves, M., Waskom, M., Clark, D., Wong, J., Loney, F., … Ghosh, S. (2018). Nipype. Software. 10.5281/zenodo.596855

Greve, D. N., & Fischl, B. (2009). Accurate and robust brain image alignment using boundary-based registration. NeuroImage, 48(1), 63–72. 10.1016/J.NEUROIMAGE.2009.06.060

Harvey, A. K., Pattinson, K. T. S., Brooks, J. C. W., Mayhew, S. D., Jenkinson, M., & Wise, R. G. (2008). Brainstem functional magnetic resonance imaging: Disentangling signal from physiological noise. Journal of Magnetic Resonance Imaging, 28(6), 1337–1344. 10.1002/JMRI.21623

Henderson, L. A., & Macefield, V. G. (2013). Functional imaging of the human brainstem during somatosensory input and autonomic output. Frontiers in Human Neuroscience, 0(SEP), 569. 10.3389/FNHUM.2013.00569/BIBTEX

Huotari, N., Raitamaa, L., Helakari, H., Kananen, J., Raatikainen, V., Rasila, A., Tuovinen, T., Kantola, J., Borchardt, V., Kiviniemi, V. J., & Korhonen, V. O. (2019). Sampling rate effects on resting state fMRI metrics. Frontiers in Neuroscience, 13(APR), 279. 10.3389/FNINS.2019.00279/FULL

Hutton, C., Josephs, O., Stadler, J., Featherstone, E., Reid, A., Speck, O., Bernarding, J., & Weiskopf, N. (2011). The impact of physiological noise correction on fMRI at 7T. NeuroImage, 57(1), 101–112. 10.1016/J.NEUROIMAGE.2011.04.018

Iyriboz, Y., Powers, S., Morrow, J., Ayers, D., & Landry, G. (1991). Accuracy of pulse oximeters in estimating heart rate at rest and during exercise. British Journal of Sports Medicine, 25(3), 162. 10.1136/BJSM.25.3.162

Jenkinson, M., Bannister, P., Brady, M., & Smith, S. (2002). Improved Optimization for the Robust and Accurate Linear Registration and Motion Correction of Brain Images. NeuroImage, 17(2), 825–841. 10.1006/nimg.2002.1132

Kelly, C., Biswal, B. B., Craddock, R. C., Castellanos, F. X., & Milham, M. P. (2012). Characterizing variation in the functional connectome: promise and pitfalls. Trends in Cognitive Sciences, 16(3), 181–188. 10.1016/J.TICS.2012.02.001

Khan, M., Pretty, C. G., Amies, A. C., Elliott, R., Shaw, G. M., & Chase, J. G. (2015). Investigating the Effects of Temperature on Photoplethysmography. IFAC-PapersOnLine, 48(20), 360–365. 10.1016/J.IFACOL.2015.10.166

Klein, A., Ghosh, S. S., Bao, F. S., Giard, J., Häme, Y., Stavsky, E., Lee, N., Rossa, B., Reuter, M., Neto, E. C., & Keshavan, A. (2017). Mindboggling morphometry of human brains. PLOS Computational Biology, 13(2), e1005350. 10.1371/journal.pcbi.1005350

Kollmeier, J. M., Gürbüz-Reiss, L., Sahoo, P., Badura, S., Ellebracht, B., Keck, M., Gärtner, J., Ludwig, H. C., Frahm, J., & Dreha-Kulaczewski, S. (2022). Deep breathing couples CSF and venous flow dynamics. Scientific Reports, 12(1). 10.1038/S41598-022-06361-X

Krause, F., Benjamins, C., Eck, J., Lührs, M., van Hoof, R., & Goebel, R. (2019). Active head motion reduction in magnetic resonance imaging using tactile feedback. Human Brain Mapping, 40(14), 4026–4037. 10.1002/HBM.24683

Krüger, G., & Glover, G. H. (2001). Physiological noise in oxygenation-sensitive magnetic resonance imaging. Magnetic Resonance in Medicine, 46(4), 631–637. 10.1002/MRM.1240

Liu, K. Y., Marijatta, F., Hämmerer, D., Acosta-Cabronero, J., Düzel, E., & Howard, R. J. (2017). Magnetic resonance imaging of the human locus coeruleus: A systematic review. Neuroscience & Biobehavioral Reviews, 83, 325–355. 10.1016/J.NEUBIOREV.2017.10.023

Liu, T. T. (2016). Noise contributions to the fMRI signal: An overview. NeuroImage, 143, 141–151. 10.1016/J.NEUROIMAGE.2016.09.008

Murphy, K., Birn, R. M., & Bandettini, P. A. (2013). Resting-state fMRI confounds and cleanup. NeuroImage, 80, 349–359. 10.1016/J.NEUROIMAGE.2013.04.001

Muschelli, J., Nebel, M. B., Caffo, B. S., Barber, A. D., Pekar, J. J., & Mostofsky, S. H. (2014). Reduction of motion-related artifacts in resting state fMRI using aCompCor. NeuroImage, 96, 22–35. 10.1016/J.NEUROIMAGE.2014.03.028

Patriat, R., Reynolds, R. C., & Birn, R. M. (2017). An improved model of motion-related signal changes in fMRI. NeuroImage, 144, *Part A*, 74–82. 10.1016/j.neuroimage.2016.08.051

Power, J. D., Mitra, A., Laumann, T. O., Snyder, A. Z., Schlaggar, B. L., & Petersen, S. E. (2014). Methods to detect, characterize, and remove motion artifact in resting state fMRI. NeuroImage, 84(Supplement C), 320–341. 10.1016/j.neuroimage.2013.08.048

Pruim, R. H. R., Mennes, M., Buitelaar, J. K., & Beckmann, C. F. (2015). Evaluation of ICA-AROMA and alternative strategies for motion artifact removal in resting state fMRI. NeuroImage, 112, 278–287. 10.1016/J.NEUROIMAGE.2015.02.063

Pruim, R. H. R., Mennes, M., van Rooij, D., Llera, A., Buitelaar, J. K., & Beckmann, C. F. (2015). ICA-AROMA: A robust ICA-based strategy for removing motion artifacts from fMRI data. NeuroImage, 112(Supplement C), 267–277. 10.1016/j.neuroimage.2015.02.064

Raj, D., Anderson, A. W., & Gore, J. C. (2001). Respiratory effects in human functional magnetic resonance imaging due to bulk susceptibility changes. Physics in Medicine and Biology, 46(12), 3331–3340. 10.1088/0031-9155/46/12/318

Salimi-Khorshidi, G., Douaud, G., Beckmann, C. F., Glasser, M. F., Griffanti, L., & Smith, S. M. (2014). Automatic denoising of functional MRI data: combining independent component analysis and hierarchical fusion of classifiers. NeuroImage, 90, 449–468. 10.1016/J.NEUROIMAGE.2013.11.046

Sasaki, M., Shibata, E., Tohyama, K., Takahashi, J., Otsuka, K., Tsuchiya, K., Takahashi, S., Ehara, S., Terayama, Y., & Sakai, A. (2006). Neuromelanin magnetic resonance imaging of locus ceruleus and substantia nigra in Parkinson’s disease. Neuroreport, 17(11), 1215– 1218. 10.1097/01.WNR.0000227984.84927.A7

Satterthwaite, T. D., Elliott, M. A., Gerraty, R. T., Ruparel, K., Loughead, J., Calkins, M. E., Eickhoff, S. B., Hakonarson, H., Gur, R. C., Gur, R. E., & Wolf, D. H. (2013). An improved framework for confound regression and filtering for control of motion artifact in the preprocessing of resting-state functional connectivity data. NeuroImage, 64(1), 240–256. 10.1016/j.neuroimage.2012.08.052

Schwarz, L. A., & Luo, L. (2015). Organization of the Locus Coeruleus-Norepinephrine System. Current Biology, 25(21), R1051–R1056. 10.1016/J.CUB.2015.09.039

Shmueli, K., van Gelderen, P., de Zwart, J. A., Horovitz, S. G., Fukunaga, M., Jansma, J. M., & Duyn, J. H. (2007). Low-frequency fluctuations in the cardiac rate as a source of variance in the resting-state fMRI BOLD signal. NeuroImage, 38(2), 306–320. 10.1016/J.NEUROIMAGE.2007.07.037

Singh, K., García-Gomar, M. G., Cauzzo, S., Staab, J. P., Indovina, I., & Bianciardi, M. (2022). Structural connectivity of autonomic, pain, limbic, and sensory brainstem nuclei in living humans based on 7 Tesla and 3 Tesla MRI. Human Brain Mapping, 43(10), 3086–3112. 10.1002/HBM.25836

Tona, K.-D., Keuken, M. C., de Rover, M., Lakke, E., Forstmann, B. U., Nieuwenhuis, S., & van Osch, M. J. P. (2017). In vivo visualization of the locus coeruleus in humans: quantifying the test?retest reliability. Brain Structure and Function. 10.1007/s00429-017-1464-5

Triantafyllou, C., Hoge, R. D., Krueger, G., Wiggins, C. J., Potthast, A., Wiggins, G. C., & Wald, L. L. (2005). Comparison of physiological noise at 1.5 T, 3 T and 7 T and optimization of fMRI acquisition parameters. NeuroImage, 26(1), 243–250. 10.1016/J.NEUROIMAGE.2005.01.007

Tustison, N. J., Avants, B. B., Cook, P. A., Zheng, Y., Egan, A., Yushkevich, P. A., & Gee, J. C. (2010). N4ITK: Improved N3 Bias Correction. IEEE Transactions on Medical Imaging, 29(6), 1310–1320. 10.1109/TMI.2010.2046908

Van Buuren, M., Gladwin, T. E., Zandbelt, B. B., Van Den Heuvel, M., Ramsey, N. F., Kahn, R. S., & Vink, M. (2009). Cardiorespiratory effects on default-mode network activity as measured with fMRI. Human Brain Mapping, 30(9), 3031–3042. 10.1002/HBM.20729

Van de Moortele, P. F., Pfeuffer, J., Glover, G. H., Ugurbil, K., & Hu, X. (2002). Respiration-induced B0 fluctuations and their spatial distribution in the human brain at 7 Tesla. Magnetic Resonance in Medicine, 47(5), 888–895. 10.1002/MRM.10145

Van Schuerbeek, P., De Wandel, L., & Baeken, C. (2022). The optimized combination of aCompCor and ICA-AROMA to reduce motion and physiologic noise in task fMRI data. Biomedical Physics & Engineering Express, 8(5). 10.1088/2057-1976/AC63F0

Windischberger, C., Langenberger, H., Sycha, T., Tschernko, E. M., Fuchsjäger-Mayerl, G., Schmetterer, L., & Moser, E. (2002). On the origin of respiratory artifacts in BOLD-EPI of the human brain. Magnetic Resonance Imaging, 20(8), 575–582. 10.1016/S0730-725X(02)00563-5

Wise, R. G., Ide, K., Poulin, M. J., & Tracey, I. (2004). Resting fluctuations in arterial carbon dioxide induce significant low frequency variations in BOLD signal. NeuroImage, 21(4), 1652–1664. 10.1016/J.NEUROIMAGE.2003.11.025

Zhang, Y., Brady, M., & Smith, S. (2001). Segmentation of brain MR images through a hidden Markov random field model and the expectation-maximization algorithm. IEEE Transactions on Medical Imaging, 20(1), 45–57. 10.1109/42.906424

Zuo, X. N., Biswal, B. B., & Poldrack, R. A. (2019). Editorial: Reliability and reproducibility in functional connectomics. Frontiers in Neuroscience, 13(FEB), 117. 10.3389/FNINS.2019.00117/BIBTEX

