## Appendix C for "Physiological Noise Correction in Brainstem Imaging: An Empirical Evaluation of fMRI Data-Driven and Peripheral Physiological Recording-Driven Methods"

### tSNR improvement due to RETROICOR over uncleaned data

As can be seen in Figure C1, variance explained by RETROICOR – in the absence of other noise regression methods – shows high overall variability across subjects. However, a general hallmark activation pattern is visible around the Circle of Willis, third, and fourth ventricle as well primarily the basilar artery. Explained variance resembles Circle of Willis structural patterns across subjects, with only a few subjects (010 and 012) showing more unspecific spatial patterns of explained variance. Note that subject 012 has a high rate of interpolated cardiac data (see Figure B1), which illustrates the potential consequences that degraded cardiac data quality can have on physiological nuisance regression. However, as can be seen in subjects 006 and 024, a reduced amount of data quality and subsequent interpolation does not necessarily mean a complete failure of RETROICOR across the run. Furthermore, it has to be noted that specificity also strongly varies between subjects. While subjects such as 002 or 014, show patterns restricted to areas most sensitive to cardiac pulsatility (Kim et al., 2021), subjects such as 001 or 029 show many wider-spread spatial patterns of explained variance. It has to be kept in mind that RETROICOR is a combination of cardiac and respiratory data and thus differing overall impacts of either might hint towards different patterns. While the magnitude of explained variance differs between subjects, the spatial pattern of peaks remains consistent. Given the direct external measurement of the cardiac and respiratory cycles, it can be expected that without severe reduction of data quality, its' explanatory value is retained independent of run length or TR.

### tSNR improvement due to ICA-AROMA over uncleaned data

Evaluating the variance explained by AROMA regressors in Figure C2, we can see a higher and less pattern-specific overall amount of explained variance. The noise component selection criteria are not originally intended for physiological noise correction. As deduced from group tSNR results, looking at the spatial distribution of the correction effect, it is visible that nonetheless there is a consistent impact of AROMA correction around the Circle of Willis, the fourth and third ventricle as well as the basilar artery (subjects 005, 008, 017, and 030). Note, as no relative comparison to other methods was made here, a lacking visual pattern of areas highly impacted by physiological noise in other subjects, can be the result of generally higher amounts of explained variance across the brain. As one of the noise component selection criteria is edge artifacts with CSF masks, the area around the 4th ventricle being most affected by cardiac pulsatility shows a high impact of cleaning. It stands out that in a majority of subjects (e.g. subject 1) a high amount of variance is explained surrounding the most inferior part of the included brainstem, traditionally extremely highly impacted by magnetization artifacts

(Brooks et al., 2013). However, a spatial pattern similar to RETROICOR is not consistently discernible.

tSNR improvement due to aCompCor over uncleaned data

Figure C3 shows the variance explained as a consequence of performing aCompCor nuisance regression. While generally, the overall level of explained variance is lower, patterns are emerging in this overview different from AROMA. As visible in subjects 004, 010, 012, 022, 024, 025, and 027, aCompCor is sensitive to striping artifacts, which in this case most likely arise from the used multi-band acquisition method with a factor of three. Similar to RETROICOR however, it also shows patterns of Circle of Willis variance in subjects 009, 013, 014, 016, and 028, possibly competing as expected with RETROICOR-based nuisance regression.

Figure C.1

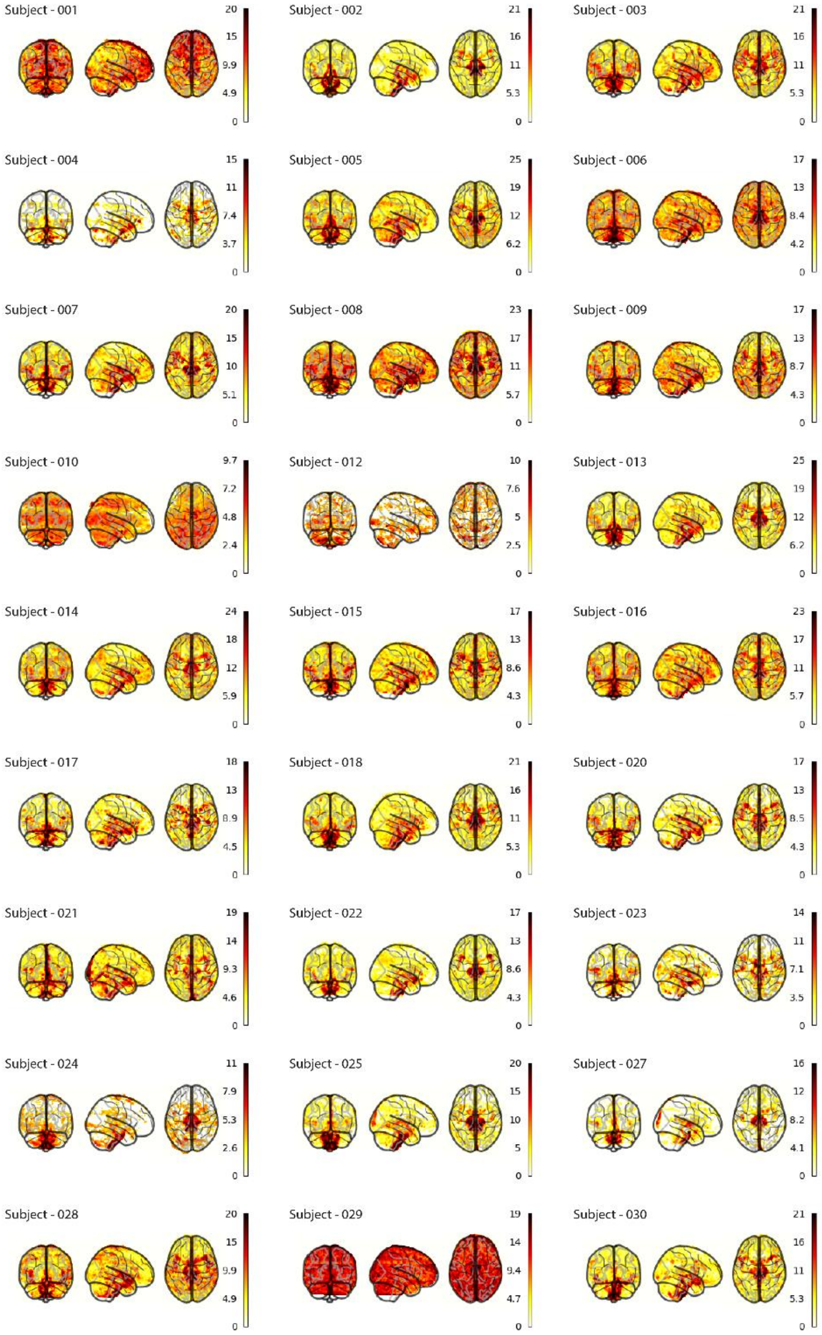


Figure C.1. Subject-wise spatial representation of individually explained variance attributed to RETROICOR (thresholded at $p_{FDR} < .05$).

Figure C.2

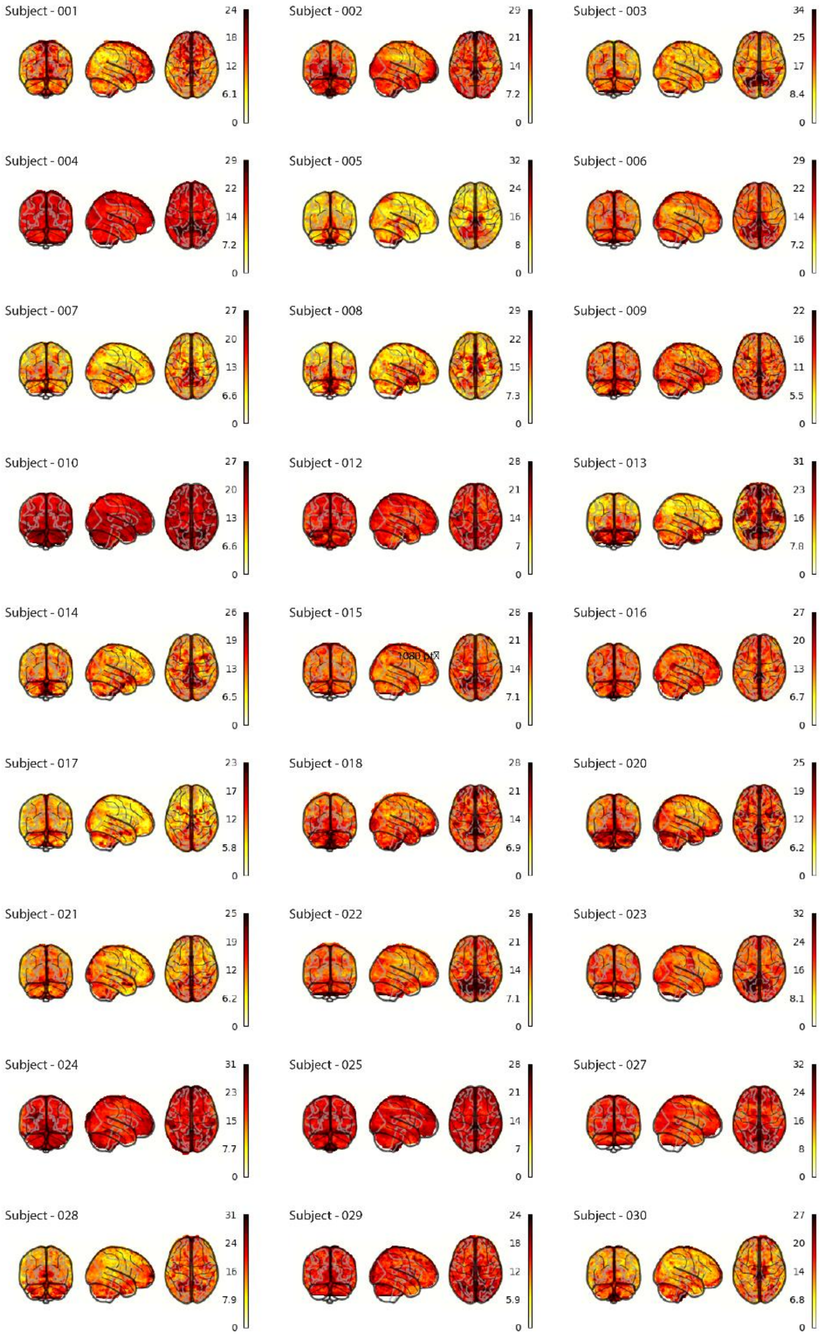


Figure C.2. Subject-wise spatial representation of individually explained variance attributed to ICA-AROMA (thresholded at $p_{FDR} < .05$).

Figure C.3

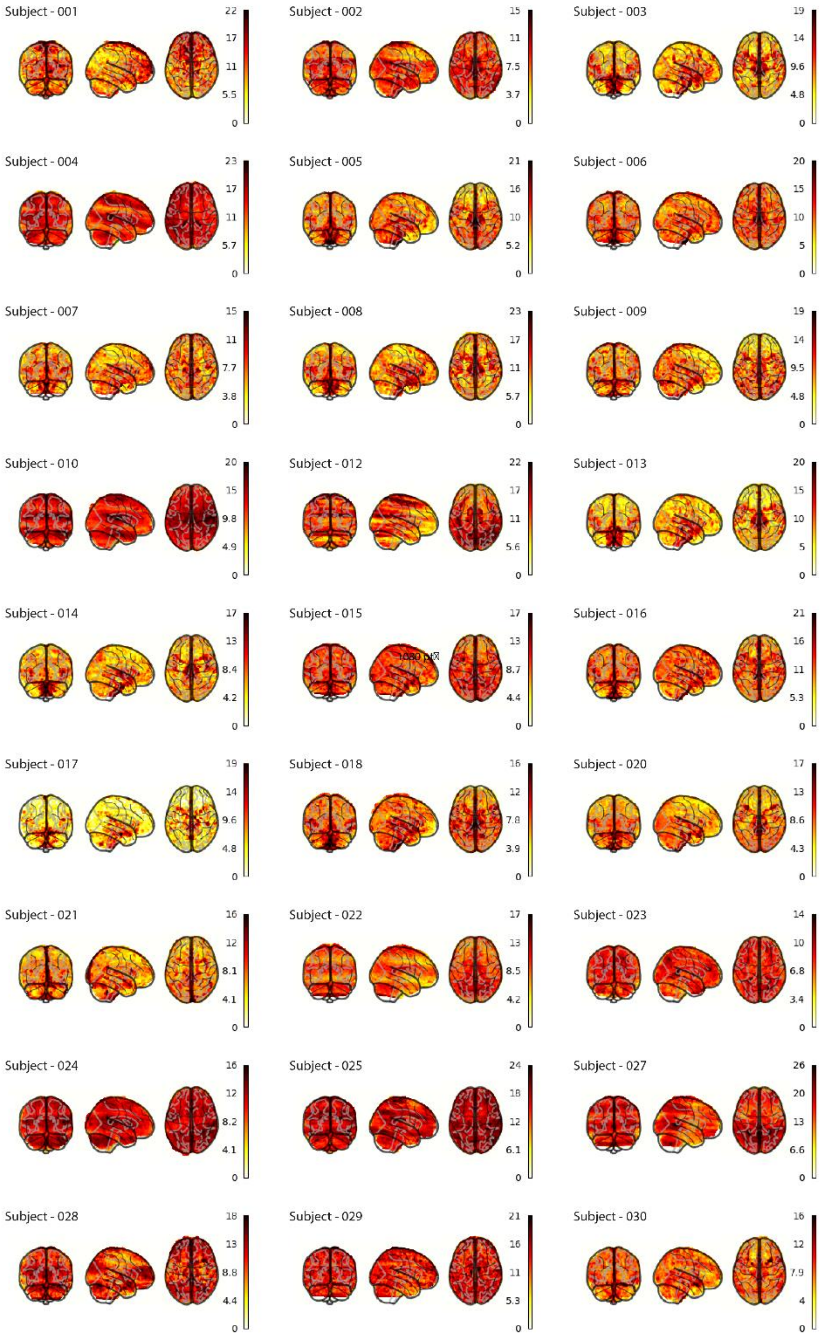


Figure C.3. Subject-wise spatial representation of individually explained variance attributed to aCompCor (thresholded at $p_{FDR}$ <.05).

Figure C.4

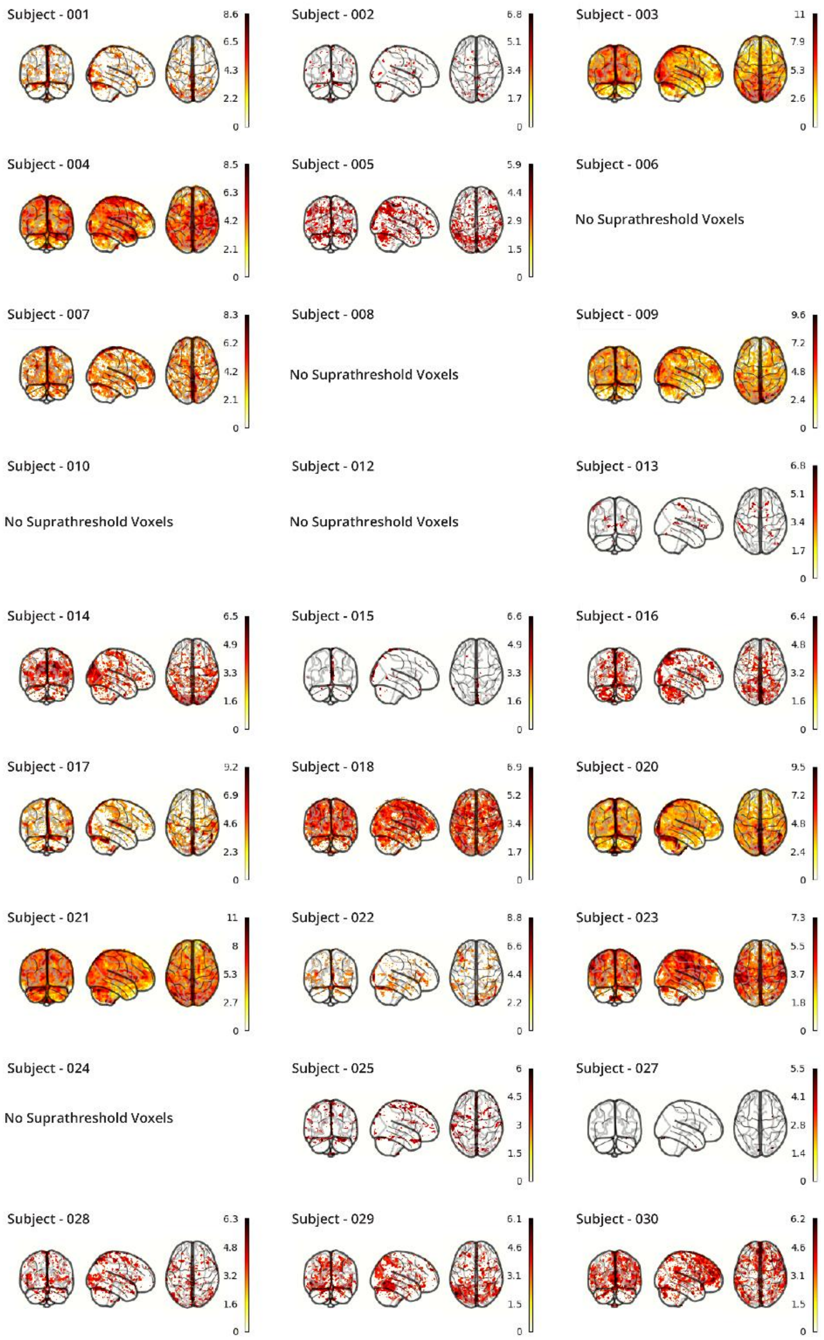


Figure C.4. Subject-wise spatial representation of individually explained variance attributed to heart rate (thresholded at $p_{FDR}$ <.05).

Figure C.5

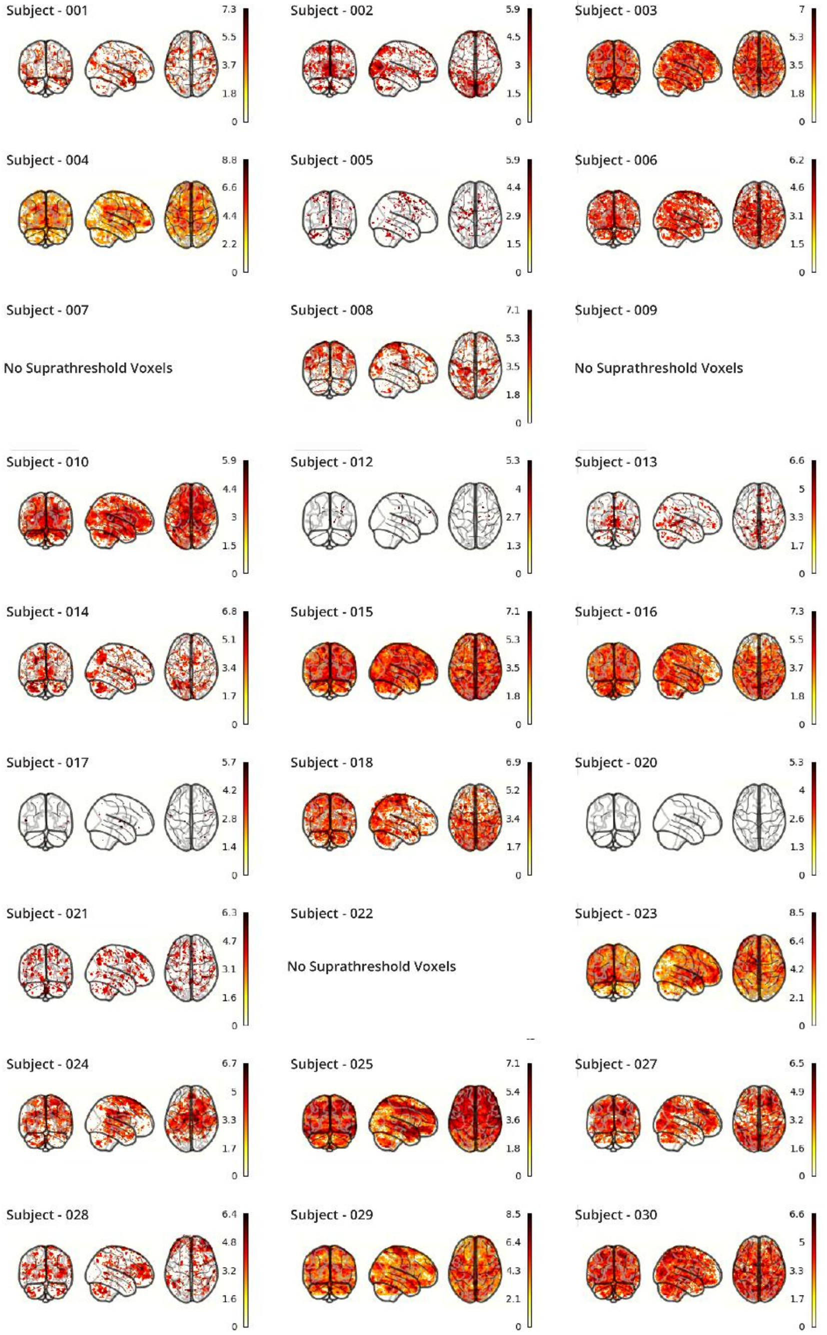


Figure C.5. Subject-wise spatial representation of individually explained variance attributed to respiratory volume per unit time (thresholded at $p_{FDR} < .05$).
