## Appendix A for "Physiological Noise Correction in Brainstem Imaging: An Empirical Evaluation of fMRI Data-Driven and Peripheral Physiological Recording-Driven Methods"

While T1-TSE allow for high in-plane resolution, they are limited by the relative slice thickness for a structure only 14.5mm in length (German et al., 1988). The positioning of the field of view thus strongly influences the possibility of detecting a three-dimensional structure with a slice thickness of about 3mm (Tona et al., 2017). As compared by K. Y. Liu and colleagues (2017), most studies to that date used higher slice-thickness to account for possible distortions, while taking advantage of the elongated shape of the LC. To increase the z-dimension resolution for better ROI segmentation and smooth slice transitions for easier manual segmentation, we deployed an in-house super-sampling approach of combining two T1-TSE sequences in a shifted z-dimension position of 1.5mm.

In line with the common practice, we acquired 3-mm thick slices with a FOV perpendicular to the brain stem at the floor of the fourth ventricle (Liu et al., 2017). Prior attempts showed that averaging three runs led to sufficient in-plane SNR. To implement the super-sampling procedure, we completed six total runs, three in the original FOV position and three with shifted z-coordinates (1.5 mm), while maintaining the transverse plain parameters. These original- and shifted runs were run alternately, changing the FOW z-coordinates by respective subtraction and addition of 1.5mm.

To combine the TSE runs collected in different orientations, three new image matrices were created with 32-rows (slices) and filled in an interleaved fashion from the respective original and shifted pair of consecutive runs (1&2, 3&4, 5&6). Subsequently, the original six TSE images were registered to the newly created averages interleaved image as a target, using mutual information for interpolation of the resampled resolution from 16 to 32 slices. By following this approach, we both tried to account for movement in between TSE-runs as well as for the partially shared information between the original and shifted sequence, creating a higher resolution – albeit smoothed – image for better detection of LC on a slice-by-slice level. As can be seen in Figure A1, this procedure enhanced visibility and allowed for easier determination of upper and lower boundaries. Purposefully, the resulting 1.5mm slice thickness corresponds to the isotropic resolution of our functional sequence, allowing for easier alignment and transformation.

Since the combined-TSE sequences only contained slice information from think 3mm slices, registration with standard methods is classically difficult. Thus, the T1-weighted images were resampled into TSE-resolution to subsequently register the TSE-combination to the T1-weighted image. As a consequence, sufficient registration to T1 could be performed to align the TSE-combination for segmentation and consequent resampling into functional EPI resolution.

To create a functional mask using this approach we manually segmented the resulting image by starting at the highest intensity voxel identified as locus coeruleus bilaterally and then moving (superior/inferior) along the Z-dimension. As the chosen approach led to substantial smoothing, we opted for the sake of simplicity and reproducibility to overlay a 1.5mm sphere on the highest intensity voxel on each slice. All subject masks were created on the image of the structural pre-processing pipeline in orientation with the existing T1-weighted image.

Using this approach, for each participant, we created a non-binary LC mask which was then used for weighted extraction of the signal by resampled into EPI resolution.

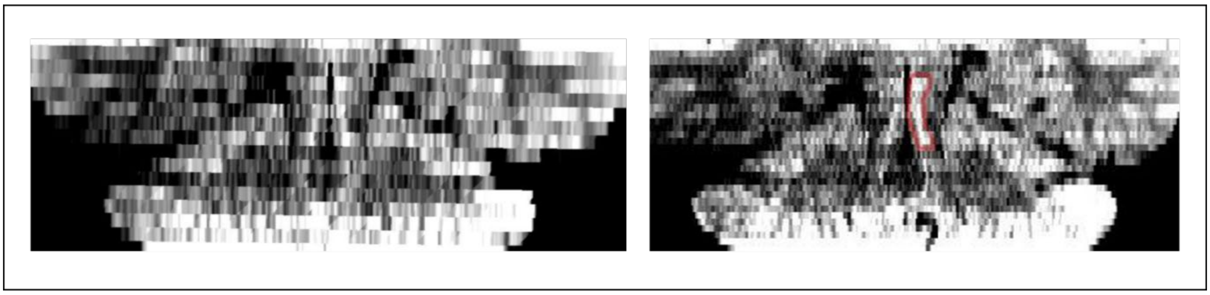

Figure A1. Original 16 slices of TSE-sequence visualizing the brain stem (left), and recombined and super-sampled TSE-sequence visualizing the brainstem with outlined LC (right).
