## Appendix B for "Physiological Noise Correction in Brainstem Imaging: An Empirical Evaluation of fMRI Data-Driven and Peripheral Physiological Recording-Driven Methods"

Cardiac processing for RETROICOR

The data quality of physiological recordings is crucial to the calculation of meaningful physiological nuisance regressors via RETROICOR and commonly used finger pulse sensors – in difference to ECG – are sensitive to motion and body temperature (Iyriboz et al., 1991; Khan et al., 2015). To assess subject variability, we calculated the amount of unusable heart rate data for nuisance regression per subject. Here, this includes a variable interval of rejection between four to eight seconds (mean = 5.31) at the beginning of the scan interval due to scan initialization. As can be seen in Figure 2, some participants displayed low quality of cardiac data. As only partial second-level analysis was performed, these subjects will still partially be used to exemplify the specificity of a RETROICOR effect. To produce full regressors, missing data were interpolated using average HR frequency of pre-post rejection intervals as implemented using an in-house pipeline.

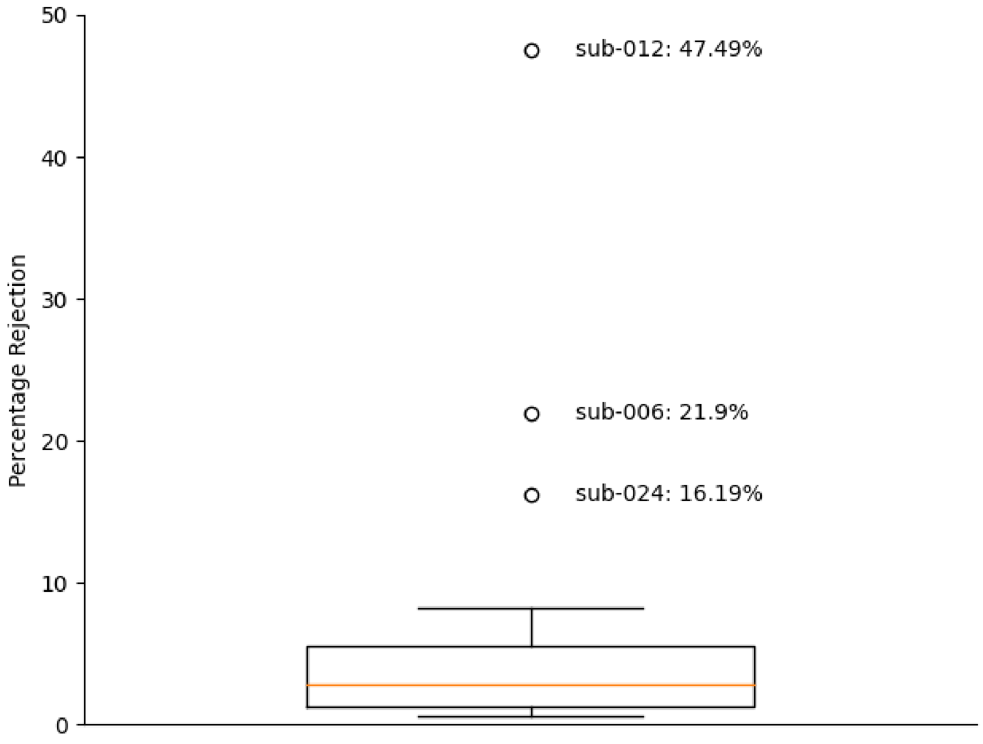


Figure B1. Percent rejection rate of raw heart rate data. Outliers described by an individual lying outside of three times inter-quartile range (IQR). Bars represent 75% IQR.
